# Cell cycle controls pathogenic processes and mycotoxin production in *Fusarium graminearum*

**DOI:** 10.1101/2025.05.01.651613

**Authors:** Thomas Svoboda, Csanad Zsiros, Florian Kastner, Michael Sulyok, Joseph Strauss

**Affiliations:** Institute of Microbial Genetics, Department of Agricultural Sciences, BOKU University Vienna, Campus Tulln, Konrad Lorenz Strasse 24, 3430, Tulln an der Donau, Austria; Institute of Bioanalytics and Agro-Metabolomics Department of Agrobiotechnology (IFA-Tulln) University of Natural Resources and Life Sciences, Vienna, Konrad Lorenz Straße 20, 3430 Tulln, Austria

## Abstract

RAS proteins control the cell cycle in all eukaryotes and lead to cancer in mammals when mutated to permanent activity. We previously isolated a spontaneous mutant of the major cereal pathogen *Fusarium graminearum* with a permanently active RAS allele due to a mutation in the nucleotide exchange factor GAP that is needed to inactivate RAS (Ras-GAP). In this study we evaluate the impact of a Ras-GAP deletion in F. *graminearum* on the phenotype and the transcriptomes of the pathogen and the host plant during infection. The mutant showed an altered secondary metabolite profile and significantly reduced virulence on wheat. The associated fungal transcriptome revealed that the mutant is unable to enter the pathogenic state and consequently, mutant cells do not switch from a saprophytic to a pathogenic program. While the wild type reprogrammed the expression of 953 genes during this switch, only six genes were significantly changed in the mutant. Genes most affected are involved in cell cycle control, response to nitrogen limitation and pathogenesis. Also, the plant responded differently and only mildly to the presence of the continuously proliferating, but avirulent fungal strain. Our data for the first time demonstrate that downregulation of the cell cycle is necessary for the production of virulence factors and pathogenicity in *F. graminearum*.

## Introduction

*Fusarium graminearum* is a plant pathogenic fungus which is able to infect different cereal crop plants resulting in reduced yield as well as mycotoxin contamination. Many studies including transcriptomics have been conducted to shed light into the genes and mechanisms involved in virulence of the fungus ^1^. While *Colletotrichum species* and *Magnaporthe oryzae* form appressoria which are emerging from a single melanized cell ^2,3^, *F. graminearum* forms appressoria like structures called infection cushions. Similar infection structures are also formed by *Botrytis cinerea, Sclerotinia sclerotiorum* and *Rhizoctonia solani* ^4–6^. Upon wheat infection 140 carbohydrate active enzymes (CAZymes), 80 putative effectors (PE) as well as 12 secondary metabolite gene clusters are upregulated in *F. graminearum* with 248 genes specifically upregulated only in the infection cushions ^7^. In *Ustilago maydis*, it has been demonstrated that a pheromone induced G2 cell cycle arrest is essential for successful corn infection. Pheromone recognition inhibits Kap123, the importin of the phosphatase Cdc25, which is needed for the G2 cell cycle progression ^8^. Similarly, in response to surface signals, appressorium formation in *Magnaporthe oryzae* is regulated by the Pmk1 MAP kinase pathway. To further develop a functionally competent appressorium, regulated cell death is required that also depends on cell cycle checkpoints^9^.

A crucial factor involved in regulation of the cell cycle is Ras which, in its active form, triggers downstream MAP-kinase signaling and activates proliferation-related genes ^10^. The conversion from the inactive to the active form is of Ras is regulated by nucleotide exchange factors (Ras-GEF) that load activating GTP onto Ras, whereas the switch back to the inactive form is triggered by a GTP to GDP conversion through activation of the intrinsic Ras-GTPase activity by Ras-GTPase activating proteins (Ras-GAP). Disruption of one of the two Ras-GAP paralogs (IRA1 or IRA2) in *Saccharomyces cerevisiae* results in suppression of lethality caused by Cdc25 mutation as well as sensitivity to heat shock and nitrogen starvation ^11,12^. For different fungal pathogens the impact of Ras and GAPs in pathogenicity has been investigated to some extent. In *Colletotrichum obiculare* disruption of the unique GAP CoIRA1 resulted in abnormal infection related morphogenesis concomitant with an increase of intracellular cAMP levels comparable to permanently active CoRAS2 strains ^13^. Four different Ras-GAPs have been characterized in *Ustilaginoidea virens* among which UvGAP1 is crucial for development and pathogenicity ^14^. In *F. graminearum* RAS2 is indispensable for full virulence. RAS2 has initially been reported to play a minor role in cAMP signaling, while it plays an important role in phosphorylation of Gpmk1, a MAP kinase involved in, and expression of FGL1, a secreted lipase ^15^. A later study showed that the guanine exchange factor RasGEF (FgCdc25) directly interacts with Ras2 proposing that Cdc25 is responsible for the modulation of cAMP and MAPK signaling pathways, fungal development and DON production ^16^. Disruption of Ras2 in *Fusarium circinatum* resulted in delayed conidial germination, smaller colonies reduced lesion size but a higher number of macroconidia ^17^.

In a previous study where the genomes of four phenotypically different *Fusarium graminearum*, PH-1, strains were compared, we identified one mutation in a Ras-GTPase activating protein in one of the strains (Svoboda et al., manuscript in preparation). The mutation is located in FGSG_00355 where the codon of amino acid 72, arginine (CGA) was exchanged to TGA, a stop codon (FGSG_00355^R72*^) resulting in loss of function. FGSG_00355 has a high similarity to iq-containing GTPases and Ras-GTPases indicating a role in the cell cycle. We employed a FGSG_00355^R72*^ Ras-GAP mutant strain and observed altered cell cycling, developmental changes, significant reduction of virulence despite strong proliferation and mycelial growth. We analyzed the fungal and the plant host transcriptomes during different infection stages and found an altered fungal transcriptome during early infection and a strongly altered defense response of the plant challenged with the Ras-Gap knock out strain.

## Results

### FGSG_00355 is highly similar to other fungal Ras-Gap proteins

A multiple sequence alignment with described Ras-GAPs of *Saccharomyces cerevisiae, Aspergillus nidulans* and *Magnaporthe oryzae* showed that the Ras-GAPs of Aspergillus, *Magnaporthe* and Fusarium are shorter than IRA1 and IRA2 of *S. cerevisiae* (Supplementary file 1). AN4998, FGSG_00355 and MoSMO1 show a high conservation among each other while deviating more from the ScIRA1 and ScIRA2. The phylogenetic tree indicated that FGSG_00355 is closer related to MoSMO1 than to the other aligned genes.

To confirm that this mutation was indeed responsible for the phenotype we reproduced the C → T mutation in FGSG_00355^R72*^ at position 214 in a PH-1 wild type strain. Already from phenotypic appearance of the emerging transformants we were able to distinguish correct transformants from the rest due to the faster growth. Two correct transformants were confirmed by PCR screening (supplementary Figure 1) and the genomes were sequenced to confirm that the mutation was introduced exclusively at the desired locus and no other high-impact mutations are present in our PH-1 wild type strain.

### Mutation of FGSG_00355 results in developmental changes, altered chemical profiles and loss of virulence

The FGSG_00355^R72*^ transformants showed a radial growth velocity comparable to the wild type on minimal media and on complete media, however, only the wild type strain appeared red on minimal media after one week (Figure 1A) indicating that secondary metabolism might be affected. Similar to the spontaneous mutant isolated previously (Svoboda et al., manuscript in preparation), both FGSG_00355^R72*^ transformants presented the typical fluffy growth phenotype indicating that hyphal morphology and developmental processes are affected by the mutation. We also tested formation of perithecia, the sexual fruiting bodies, on carrot agar plates and found that the mutant strain produced less perithecia than the wild type, however pigmentation of these structures did not differ much between the wild type and the mutant (Fig. 1F).

**Figure 1:**
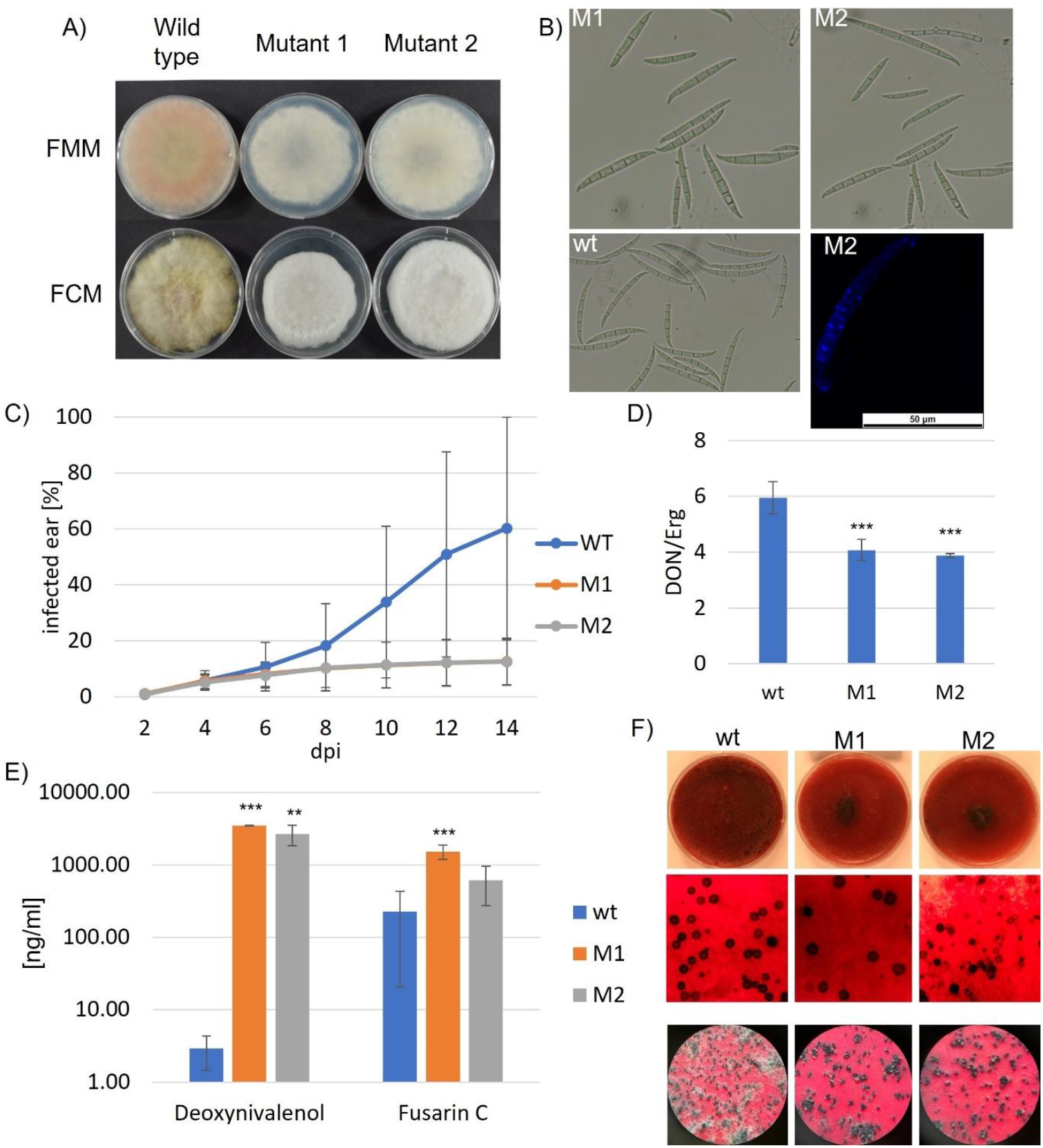
Characterization of two independent Ras-GAP mutant strains; A) growth on Fusarium minimal media (FMM) and Fusarium complete media (FCM); B) appearance of the spores under the microscope and on the bottom right a single spore of a mutant strain with nuclei stained. Multiple nuclei are visible between two septa; C) spread of wheat infection after point inoculation with the respective strain; D) DON production normalized for the fungal biomass present in the wheat ear; E) DON and Fusarin C production on YPD plates show significantly higher DON production in the mutant strains; F) Perithecia formation on carrot agar: the whole plate after 2 weeks (top), under the microscope (middle), after 1 month (bottom)

Inspection of the few conidial spores produced by strain FGSG_00355^R72*^ revealed that they are less curved and in most of them septa were missing or strongly reduced. At the same time some nuclei appear to be stuck in the separation process. NucBlue™ staining and confocal microscope analysis showed that the mutant spores carry much more nuclei per spore than the wildtype and when septa were present, multiple nuclei were visible within one septal compartment of the mutant whereas in the wild type only one nucleus per septal cell is present. Taken together, these phenotypical observations indicate that the mutant lost control over cell cycle and mitosis, a feature that is consistent with the proposed role of RAS-Gap in turning down the cell cycle by inhibiting the function of Ras (Figure 1B).

When we tested secondary metabolite profiles of the mutant strains we found a surprisingly high DON production in axenic cultures in both mutant strains as compared to the wild type (Figure 2E). The results of the virulence test also revealed that the Ras-GAP mutant strains are able infect the inoculated spikelets, however, they are hardly able to spread within the ear (Figure 2C). This was rather surprising due to the high DON production in axenic culture that serves as a major pathogenicity factor in *F. graminearum* wheat infection ^18^. Surprisingly, the analysis of the infected ears for secondary metabolites revealed that the plants infected with mutant strains contained significantly lower amounts of almost all analyzed mycotoxins (supplementary table 1). To normalize the SM levels for biomass, we measured ergosterol content in the infected material. After biomass normalization, there were still significantly lower amounts of DON (Figure 2D) present in the ears infected by the Ras-GAP mutants. And nivalenol, zearalenone and calonectrin were even below the limit of detection in both mutant strains, while being detected in the wild type.

**Figure 2:**
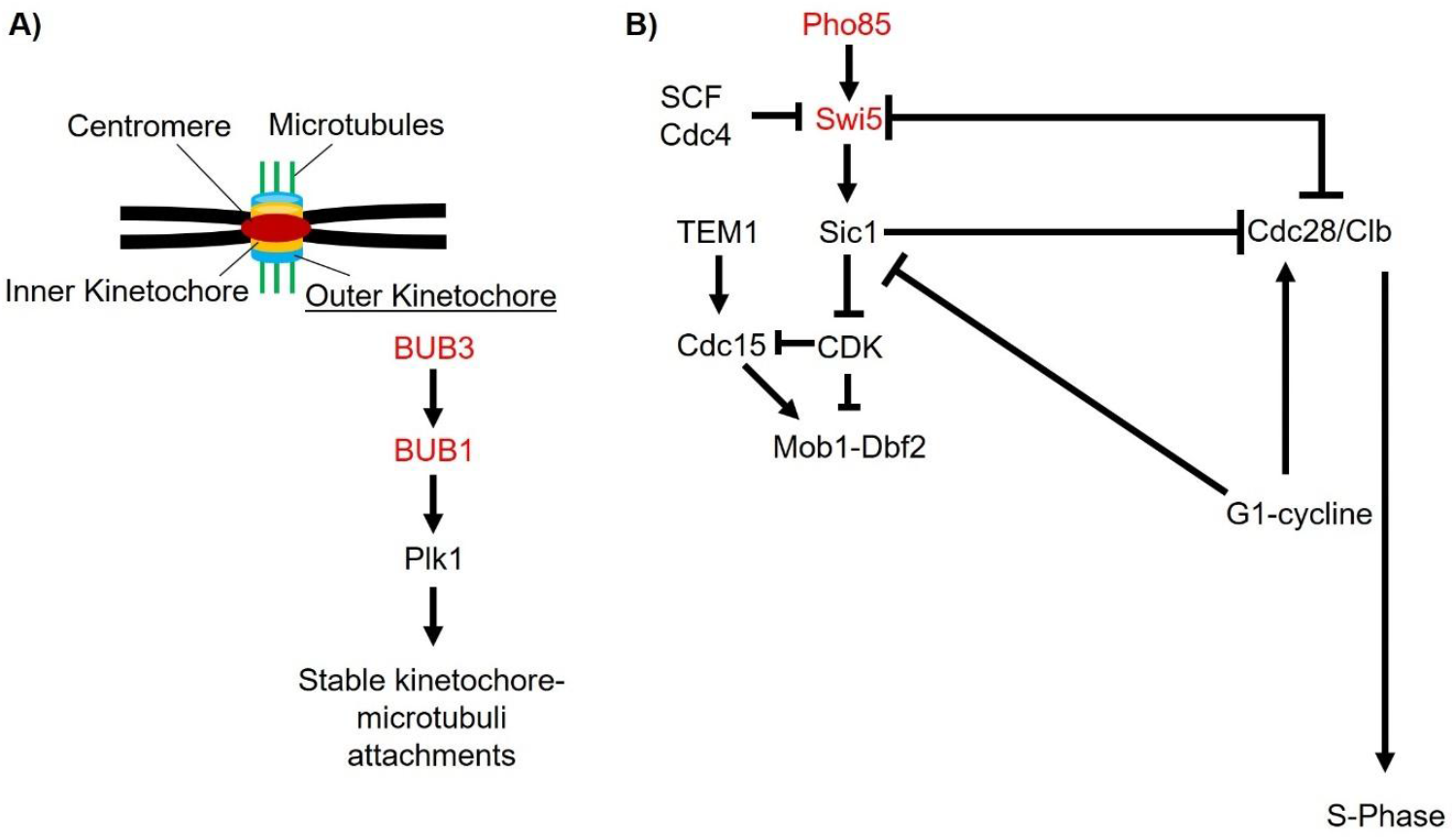
possible impact of the Ras-GAP mutation on the cell cycle during infection; A) BUB3 and BUB1 which are responsible for the control of the kinetochore-microtubule attachment are both significantly downregulated; B) Pho85 and consequently Swi5 are both significantly lower in the mutant which may have an impact on exit from mitosis but also do no longer repress Cdc28 mediated G1/S phase transition; red font indicates a significant downregulation of the homologous *F. graminearum* genes in the Ras-GAP LOF mutant

### The infection transcriptomes show strongly altered responses in the pathogen and the plant

In order to gain insights into the virulence mechanisms that changed in the Ras-GAP mutant in its host plant, we performed transcriptome analyses at two different time points after inoculating flowering wheat heads. For the transcriptomic profile, each spikelet of six ears was inoculated with a defined amount of conidia or water as control and incubated for two or four days. Upon harvesting, three ears of each treatment were pooled, RNA extracted and sequenced by Illumina Hi-seq. Hence the results of each sample represent the mean expression pattern of each gene upon infection of three different ears.

### Loss of function mutation in Ras-GAP leads to downregulation of various pathways at the early infection phase

A KEGG enrichment analysis of the different expression pattern between the wild type and the mutant strain revealed that in the mutant strain genes in three major pathways are affected by lower expression (p<0.05) during early stage (supplementary file 2): general metabolism (fgr00330, fgr00053, fgr01232, fgr00350), genetic information processing (fgr03013) and cellular processes (fgr04111). The metabolic pathways affected were tyrosine, arginine and proline metabolism, ascorbate and aldarate metabolism, and nucleotide metabolism.

Strikingly, many of the genes less expressed in the mutant are associated with functions in cell division and regulation of the cell cycle. Generally, negative regulators of the cell cycle are less expressed in the mutant while positive regulators are found at higher expression levels. For example, FGSG_07139, which is supposed to belong to the cohesin complex subunit SA-1/2, is lower expressed in the mutant by log2FC of -7.70. Also, a DNA replication licensing factor (mcm3), which is activated during the G1-S phase to unwind the DNA and maintain the DNA replication fork elongation ^19–21^ is lower expressed (FGSG_01920: log2FC -7.60). Other less expressed genes structural maintenance proteins of chromosome 1 (FGSG_01910) and chromosome 2 (FGSG_05105) also the cell cycle arrest protein BUB3 pops up which plays a crucial role in the chromosome-microtubule sensing. As long as the kinetochore-microtubule interaction is not established, Bub3 delays the entry to the anaphase ^22–25^. Bub1 (FGSG_01347), mitotic arrest deficient 1 (Mad1) in yeast, is a serine threonine kinase (log2FC -7.27) which is essential for correct chromosome alignment as well as for the spindle-assembly checkpoint signaling ^26^. Both, BUB1 and BUB3 are crucial for telomer amplification and loss of these proteins results in shortened and fragile telomers ^27^. An extended cell cycle arrest is crucial to repair potential DNA damage ^28^ which could also be impaired in the mutant strain.

The interaction of Grr1 and Skp1, which is part of SCF, plays a crucial role in the degradation of the cell cycle regulators Cln1 and Cln2 ^29^ which are involved in Cdc28 activation ^30^. Lower expression of Pho85 upstream of Swi5 would lead to less inhibition of Cdc28 and thus to an active cell cycle in the Ras-GAP mutant in contrast to infectious wild type that would stall the cell cycle at this point (references 31-35).

### Several virulence factors are significantly lower expressed in the mutant strain at the early infection phase

In *Fusarium graminearum* several virulence factors are known which are divided between carbohydrate active enzymes (CAZymes), effectors and secondary metabolite gene clusters (SMC). At the early time point, a few previously described effectors are significantly lower expressed in the Ras-GAP mutant compared to the wild type strain indicating a cell cycle dependent regulation. FGSG_10563 (log2FC = -7.73) encodes for a LysM domain containing protein (IPR018392) which is generally described to be crucial to overcome plant defense ^7,31^. The second one (FGSG_10595) belongs to the class of peptidase proteinase inhibitor I9 which generally inhibit plant defense and promote the spread of the fungus (reviewed by Jashni et al. 2015). The third protein significantly lower in the mutant strain (log2FC = -8.43) is annotated as necrosis inducing protein NPP1 (FGSG_06017) which has been described in *Phytophthora cinnamomic* to be expressed after 36 hours due to the complex attack/defense interaction between the host and the pathogen ^33^. Additionally, FgHyd5 (FGSG_01831), which plays a crucial role for initial infection ^34^, is significantly lower expressed at the early infection phase (log2FC = -8.50).

### Strong difference between wild type and Ras-GAP mutant two days post infection

The comparison of the expression pattern of the wild type and the mutant strain 2 dpi showed that at this early stage of infection 607 genes are significantly lower expressed in the mutant compared to the wild type while only 12 genes are significantly higher expressed. However, four days post inoculation there is only one gene remaining significantly lower expressed in the mutant strain compared to the wild type. A similar pattern is observed when we compare the changes in the respective strain between two and four dpi. While in the mutant strain only four genes and two genes are significantly higher and lower expressed, respectively, at the later time point a much stronger change is observed in the wild type strain where ten genes are higher but 943 genes are significantly lower expressed after 4 dpi. Among those 943 downregulated genes also Ras-GAP (FGSG_00355) is found with a log2 fold change of -6.67 (p = 0.0072) indicating a crucial role of cell cycle regulation at the early stage of infection. These changes indicate that due to the mutation of Ras-GAP the mutant is hardly able to control its metabolism in response to environmental factors. Analysis of the trichothecene biosynthesis genes revealed that FGSG_03538 (*TRI10*) is significantly lower expressed in the mutant strain (log2FC = -7.30, p = 0.0057). FGSG_03538 is a regulatory protein which controls the transcription of *TRI6* which consecutively regulates the expression of *TRI5* and *TRI4* catalyzing the initial steps of trichothecene biosynthesis (Kimura et al. 2007; Tag et al. 2001). This is also in line with the results of the secondary metabolites extracted from infected wheat normalized for fungal biomass where the mutant strains produce significantly lower amounts of DON (Fig. 1d).

Functional annotation of the differentially regulated genes in the respective strain between two and four dpi revealed that in the mutant exclusively one downregulated gene could be assigned to energy (nitrogen) metabolism (FGSG_02970). In the wild type strain genetic information processing (ribosomal RNAs) was strongly induced 4 dpi indicating that growth is forced once the infection is established. A closer look into the data revealed that FGSG_20075, which is involved ribosome biogenesis, is significantly upregulated in the wild type after 4 dpi turned out to be significantly higher expressed in the mutant strain at 2 dpi compared to the wild type. This is a strong indication that the mutant is not able to adapt to environmental changes and switch to infection mode due to the constant growth signal.

Comparison of the genes deregulated after four dpi taking two dpi in the respective strain into account revealed that among the significantly changing genes there was no overlap between wild type and mutant strain (supplementary Figure 2).

We further compared the expression profile of the wild type and the mutant at the different time points as well as the changes of the expression between two and four days. Enrichment analysis revealed that the large subunit of the 28S/25S rRNA is significantly higher in the mutant compared to the wild type strain 2 dpi. However, analysis of the changes in the wild type over time showed a significantly higher expression of both the small (18S) and the large ribosomal subunit after 4 dpi compared to 2 dpi indicating induction of growth once the infection is established. Ribosome biogenesis plays a crucial role in generating ribosomes and is further indispensable for cellular processes like proliferation, development, differentiation and apoptosis (reviewed by Jiao et al. 2023).

### Ras-GAP mutant triggers different defense response in wheat

To find out whether the different expression patterns in the two tested fungal strains trigger different defense responses in the plant, the transcripts of the ears were inspected as well. A first glance at the changes in expression in the ears over time upon infection with the two strains showed that there were much stronger changes occurring in the wild type treated ears compared to the mutant treated ones between two- and four-days post inoculation. While wild type treatment caused a significant change in expression of 1345 genes, inoculation with the mutant affected only 650 genes. Interestingly, only a few deregulated genes are shared upon wild type and mutant inoculation indicating that the strains trigger strongly altered responses.

**Figure 2:**
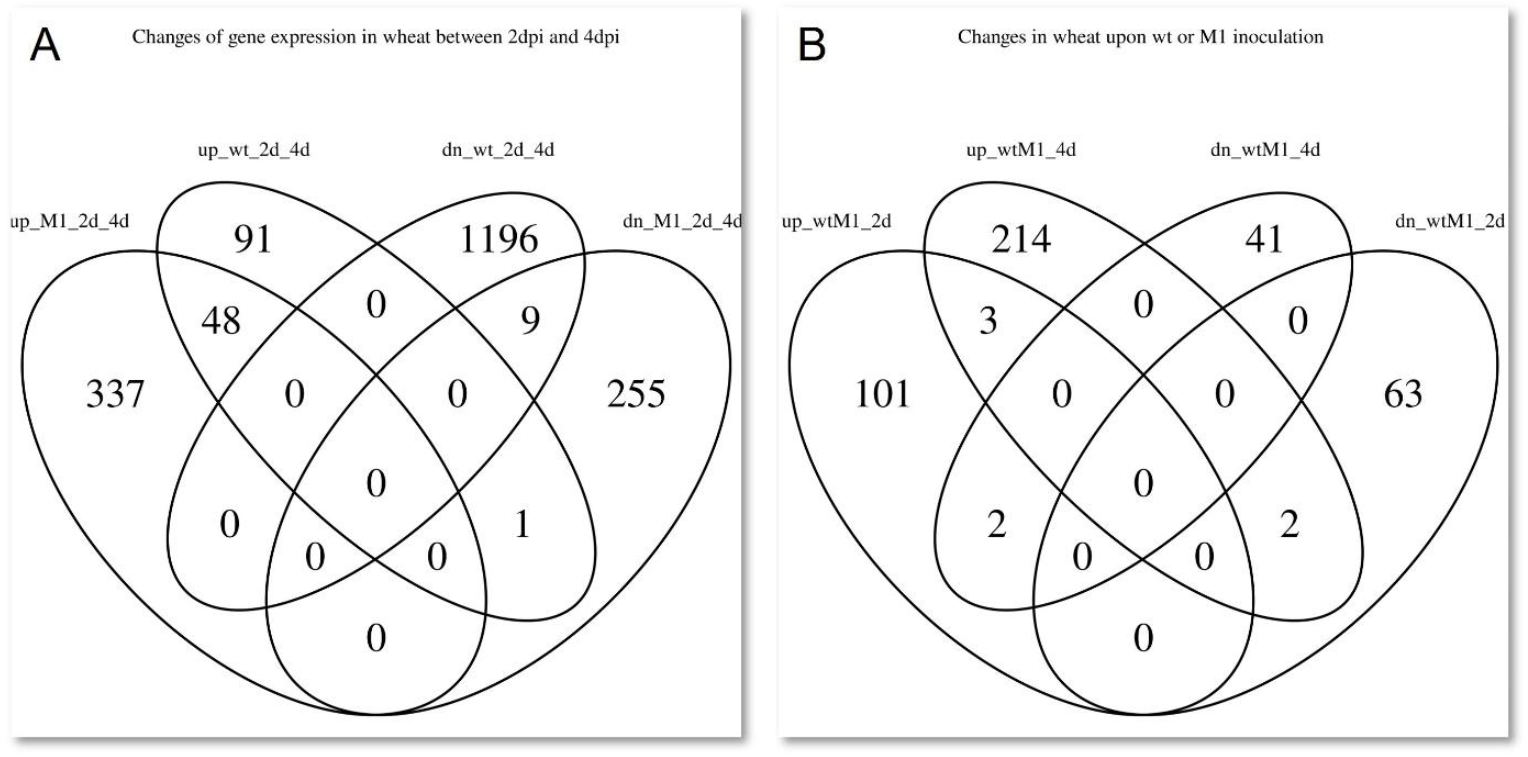
Venn diagram; A) genes which show a significant change between two and four dpi upon inoculation with the wild type or the mutant are indicated; B) genes which are significantly different expressed upon inoculation with the different strains after two and after four days are indicated

The IDs of the genes respectively shared are indicated in supplementary table 2. While nine genes were significantly downregulated between two and four dpi upon inoculation with both strains, 48 genes went significantly up. These genes seem to be generally activated in response to fungal infection. There is only one gene downregulated after inoculation with the mutant between 2 dpi and 4 dpi while it is upregulated upon wild type infection. This gene (LOC123135745) is annotated as disease resistance protein RGA2-like. Additionally, this gene is also significantly higher expressed 2 dpi upon mutant inoculation compared to wild type inoculation while after 4 days the expression is significantly lower. The second gene which is significantly higher after mutant inoculation 2 dpi but lower after 4 dpi compared to wild type inoculation is LOC123186820, a yet uncharacterized protein. On the other side, there are two genes significantly lower expressed 2 dpi but higher expressed 4 dpi upon inoculation with the mutant compared to the wild type. LOC123098147 is a DNA (cytosine-5)-methyltransferase CMT-like protein while LOC732704 is a gamma-interferon-responsive lysosomal thiol protein which has been described as a major wheat allergen ^38,39^.

Enrichment analysis of the deregulated genes showed that between two and four days after wild type inoculation, phenylalanine metabolism as well as zeatin biosynthesis are significantly downregulated in response to wild type infection while there is no gene set enrichment upon mutant inoculation. In the phenylalanine metabolism (taes00360) there are several phenylalanine ammonia lyase like genes as well as one tryptophane decarboxylase 1-like gene deregulated. Three groups of enzymes, cytokinin dehydrogenase 2, cis-zeatin-O-glycosyltransferase and UDP glycosyltransferase, are involved in the in the zeatin metabolism (taes00908) significantly higher expressed 2 dpi compared to 4 dpi after wild type treatment.

Within the differentially expressed genes shared upon inoculation with the two strains 4 dpi, there are three pathways enriched, however, only one gene involved in all three pathways is popping up. It is a chalcone synthase 2 which is involved in the flavonoid biosynthesis, the circadian rhythm of the plant as well as in tropane, piperidine and pyridine alkaloid biosynthesis. Taking the mock treated control into account, we can see that the identified gene (geneID 123051070) is already expressed 2 dpi indicating that transcription may be negatively influenced by the fungus during initial infection since after inoculation with the different Fusarium strains not a single read was detected 2 dpi. Chalcone synthases, which belong to type III polyketide synthase, are involved in plant defense as key enzyme of the salicylic acid defense pathway (reviewed by Dao, Linthorst, and Verpoorte 2011).

## Discussion

In this study we demonstrated that constant presence of Ras-GTP due to Ras-GAP inactivation causes beside an abnormal phenotype, significant changes in the transcriptome during infection. Especially proteins which are involved in the regulation of the cell cycle but also virulence factors showed a strongly altered expression in the loss of function mutant compared to the wild type. A cell cycle dependent transcriptome has been described in *Cryptococcus neoformans* and *Saccharomyces cerevisiae* by synchronization of the cell cycle to assess the periodic gene expression. It was documented that genes involved in the regulation of S-phase and mitosis were periodically transcribed in the respective phases ^41^. The regulation of S-phase is important for the formation of infection structures. In *Magnaporthe oryzae* appressoria formation is S-phase dependent and connected to the DNA damage response (DDR) pathway. In contrast, appressorium repolarization requires a DDR independent checkpoint in the S-phase ^42^. In Fu*sarium graminearum* trichothecene production in such infection structures is enhanced even though the toxin is neither necessary for infection cussion formation nor for causing primary symptoms ^43^. Analysis of the transcriptome of *F. graminearum*, revealed that especially in the symptomless wheat tissue the transcriptional inducers, *TRI6* and *TRI10*, of the TRI cluster which are responsible for trichothecene biosynthesis and full virulence ^44–46^, are increased ^47^. This is in line with our study, where also a significantly lower expression of *TRI10* was found in the Ras-GAP mutant strain which may be one factor the lower virulence is owed to.

In the S-phase, the replication licensing factor mcm3 which is part of the mcm2-7 hexamer, is significantly lower expressed compared to the wild type. However, the impact of the lower expression is not necessarily detrimental for S-phase entry. This has been demonstrated by knock down of MCM3 in HEK 293T cells where no effect on S-phase entry was observed while overexpression of MCM3 inhibits S-phase entry suggesting a negative impact on DNA replication ^48^. Already in G1 phase, loop extrusion is restricted by the MCM complex which also acts as physical barrier constraining cohesin translocation ^49^. In this context, cohesin subunits which also play a role in DNA repair as well as development and gene expression, were also found significantly lower expressed in the Ras-GAP mutant strain. During G2 phase until anaphase, where the chromatids separate, the sister chromatids are connected by cohesins ^50^. A kinetochore associated pool of cohesins plays a crucial role in withstanding the spindle pulling forces in the metaphase, thereby strengthening the centromeric sister-chromatid cohesion ^51^. DASH/Dam1, also part of the kinetochore and significantly lower expressed in the Ras-GAP mutant, is present as heterodecamer assembling into a microtubule encircling ring thereby organizing eventually also other kinetochore components and the Ndc80 complex (Ndc80c) rods ^52–54^. While Ndc80c binds to the C-terminal region of Dam1, the mitotic kinase Ipl1/Aurora B can phosphorylate serine 257, 265 and 292 releasing the contact for error correction of mis-attached kinetochores ^55^. In histone-humanized yeasts where the centromere function is disrupted which usually results in aneuploidy, DASH/Dam1 mutants display euploidy probably due to the loss of Aurora B activity which is responsible for the attachment of the mitotic spindle to the centromere ^56^. Aneuploidy can also occur due to the lack of the spindle checkpoint gene BUB3. However, aneuploidy of different chromosomes does not necessarily seem to be detrimental since it is maintained over many generations providing an advantage to strains with low chromosome segregation fidelity ^57^. BUB3 and BUB1, which are both lower expressed in the Ras-GAP mutant, are both part of the spindle assembly checkpoint (SAC) as well as Mad1, Mad2, Mad3 (BubR1), Mps1 and Aurora B. SAC is a crucial checkpoint in the transition from anaphase to metaphase (reviewed by Logarinho and Bousbaa 2008). While BUB1 and BUB3 can delay anaphase onset due to reduced tension, Mad1-3 are not involved in this regulation ^59^.

Swi5 which usually peaks at the G2/M transition but enters the nucleus in the late M phase as well as Pho85 are both significantly downregulated in the mutant strain. Swi5 is a transcription factor which is activated and stabilized by phosphorylation of a serine rich region by Pho85. As consequence, Sic1, which is highly unstable and expressed in the late M-phase ^60,61^, is transcribed which consequently results in repression of CDK ^62,63^. This CDK is responsible for repression of Cdc15 which is crucial for mitotic exit and cytokinesis ^64^. Since in the mutant strain Swi5 is significantly lower as consequently the repression of CDK may be impaired. This implies that the mutant strain there is no cell cycle halt in mitosis in metaphase where the kinetochore-microtubuli attachment should be verified by BUB1 and BUB3, however, for telophase control Sic1 is no longer activated by Swi5. Consequently, Sic1 mediated repression of CDK is impaired resulting in enhanced inhibition of Cdc15 and disturbances in mitotic exit. While the cell cycle regulation is out of control in the mutant strain, at the same time various virulence factors are significantly lower expressed compared to the wild type strain indicating a cell cycle dependent regulation.

While various cell cycle control proteins are deregulated also several virulence factors are deregulated upon loss of function of Ras-GAP. LysM, peptidase proteinase inhibitor I9, NPP1 and FgHyd5 are significantly lower expressed in the mutant strain 2 dpi compared to the wild type strain. All of these proteins have been documented to be crucial for full virulence, however, dependency of those genes on the cell cycle has not been documented so far. Mentges et al. (2020) showed that upon wheat palea inoculation, *F. graminearum* actively grows over the leaf with runner hyphae and forms stationary, multicelled appressoria (infection cussions) at different spots. It has been demonstrated that cell cycle regulation and appressorium formation are connected by induction of G2-arrest in *Ustilago maydis* ^65^ or for proper cell cycle progression from G1 to S phase in *Colletotrichum obiculare* ^66^. Eventually the deregulation of the cell cycle in *F. graminearum* does not only lead to a deregulated cell cycle but also to the lack of the feedback control within the infection cussions.

Further different metabolic pathways are obviously depending on the cycling between Ras-GTP and Ras-GDP. In the early infection phase nitrogen acquisition and adaption to nitrogen poor conditions are essential for pathogenesis. In *Fusarium oxysporum* FNR1, a conserved GATA-type zinc finger domain containing protein involved in nitrogen metabolism, is crucial for full virulence ^67^. In this context we identified AreA (FGSG_08634) significantly downregulated in the mutant strain (log2FC = - 7.15) compared to the wild type at 2 dpi, one factor contributing to reduced virulence. AreA interacts with Tri10, which is also significantly downregulated in the mutant at the same time, and is further involved in DON production by ammonium suppression and the cAMP-PKA pathway ^68^. Additionally, AreA is required for adaption to and metabolization of nonpreferred nitrogen sources ^69^. AreA is usually starvation induced ^70^, however, the mutant does not seem to get the starvation signal due to the constantly active growth signal by Ras-GTP. Even though sensitivity of *S. cerevisiae* lacking IRA1 or IRA2 to nitrogen starvation ^11,12^ this is the first indication that AreA mediated nitrogen starvation response during infection is not only triggered by the lack of nutrients but also depending on the cell cycle. Extended nitrogen starvation (25 days) in heterothallic, haploid fission yeast cells results in entering the G0 phase ^71^.

The two different genotypes which show strongly diverging transcription patten during wheat infection also trigger different responses in the plant. Exclusively in wild type infected ears phenylalanine metabolism as well as zeatin metabolism are significantly higher expressed 2 dpi compared to 4 dpi. Phenylalanine is not only used as amino acid in proteins but also metabolized by phenylalanine ammonia lyase (PAL) which plays a crucial role in defense response ^72–74^. Through the phenylpropanoid pathway, hydroxycinnamic acids are formed which are further used for the formation of hydroxycinnamic acid amides (HCAAs). Phenylalanine metabolism and formation of HCAAs is a response to DON produced by Fusarium ^75^. Due to significantly lower DON production in the mutant strain upon infection, the corresponding response in the wheat plant is not induced.

On the other side we observed significant changes in the zeatin metabolism. Zeatin is a naturally occurring cytokinin. Upon exogenous application on wheat the resistance towards *Stagonospora nodorum* is enhanced by upregulation of salicylic acid metabolism and inhibition of the ethylene signaling pathway ^76^. A growth assay on plates showed that zeatin has an inhibitory impact on *Botrytis cincerea*. Similarly cytokinins generally have an inhibitory impact on fungal growth and virulence which has been demonstrated *B. cinerea, S. rolfsii*, and *F. oxysporum f. sp. lycopersici* ^77^. We can see that the strongly altered gene expression profile in the fungus upon wheat inoculation is owed to the Ras-GAP mutation. This further results in strong changes in the defense response of the plant. Several virulence related genes are no longer produced in the mutant strain and therefore the corresponding defense response is no longer triggered.

## Conclusion

Taken together, a loss of function mutation in Ras-GAP results is strongly altered transcriptomes in the fungus as well as strong changes in wheat upon infection. The mutant strain shows a strongly deregulated cell cycle where several checkpoint proteins are significantly lower expressed compared to the wild type strain. Also, different virulence factors as well as AreA, responsible for the nitrogen starvation response, are hardly expressed upon wheat infection indicating a connection to the cell cycle. On the other side, the mutant strain triggered a strongly altered response in wheat compared to the wild type. We identified 48 core defense response genes, however, the majority of the defense related genes is triggered by various virulence factors of the fungus. For future research, on the plant side, the role of the 48 core defense genes needs to be characterized and on the fungal side there are still open questions regarding how the cell cycle has an impact on the expression of virulence factors and on AreA.

## Material and Methods

### Media

#### Fusarium minimal media (FMM)

1 g/L KH_2_PO_4_, 0.5 g/L MgSO_4_.7H_2_O, 0.5 g/L KCl, 2 g/L NaNO_3_, 30 g/L sucrose. For solid media 20 g agar were added. After autoclaving 200 μl of a trace element solution (for the preparation of 100 ml trace element solution 5 g citric acid, 5 g ZnSO_4_.6H_2_O, 1 g Fe(NH_4_)_2_(SO_4_)_2_.6H_2_O, 250 mg CuSO_4_.5H_2_O, 50 mg MnSO_4_, 50 mg H_3_BO_4_, and 50 mg Na_2_MoO_4_.2H_2_O were used) were added ^78^.

Complete medium was prepared according to Pontecorvo et al.^79^.

Mung bean soup (MBS): For 1 liter MBS, 30 g mung beans are cooked in water for 30 minutes. The media is filtered to get rid of the beans followed by autoclaving.

### Ras-GAP – plasmid design for introduction of the loss of function mutation

The plasmid was constructed by using yeast recombinational cloning (YRC) using pRS426 as vector. The fragments (F1-F5) were amplified using the primers and the templates listed in supplementary table 3. The bold nucleotides in P2 and P3 indicate the mutation introduced. and transformed in yeast together with SmaI linearized vector pRS426. After YRC, the plasmids were prepped from yeast, transformed into *E. coli* followed by mini prep and a test digest and sequencing. For the transformation the fragment was amplified using primers P1 and P10 from the previously constructed plasmid and transformed into *Fusarium graminearum* according to Twaruschek et al. (2018). The transformants were selected on 40 ppm G418 followed by transfer and screening PCRs according to supplementary table 3.

### Genome sequencing of mutants

To confirm that no other mutation was randomly introduced, the genomic DNA of the correct candidates was extracted ^81^ and sent to sequencing. The genomes were aligned to the reference genome using burrow wheeler aligner version 0.7.18. Further samtools version 1.21 was used to format, sort and index the files. For variant detection freebayes v1.3.6 was used. The genome was inspected using the integrative genome viewer.

### Characterization of the mutants

Growth tests were performed on Fusarium complete media (FCM) as well as Fusarium minimal media (FMM). 10 μl of a 1*10^6^ spore suspension were pipetted in the middle of the plate. After incubation at 20°C for seven days, the plates were evaluated. Sporulation was tested in mung bean soup. Small pieces of mycelia were inoculated in 50 ml MBS followed by incubation at 20°C, 140 rpm. The spores were harvested by filtration through glass wool tips to get rid of mycelia. After centrifugation the supernatant was discarded and the spores were suspended in 1 ml sterile H_2_O. Perithecia formation was tested on carrot agar as described by Cavinder et al. (2012).

Deoxynivalenol (DON) production was tested on complete media, minimal media as well as minimal media supplemented with ornithine. Further, the virulence was tested as well as in planta toxin production by inoculation of two spikelets (four florets) in the middle of the ear of the susceptible cultivar Apogee with 10 μl of a 4*10^5^ spore suspension. A plastic bag which was previously sprayed with water was put over the ear for 24 h to maintain humidity. The plants were incubated at 20°C with a 16h/8h day/night cycle. The progress of infection was evaluated every two days. After 14 days the samples were harvested, frozen in liquid nitrogen and homogenized using Retsch mill (f = 30 s^-1^, 30 sec). Secondary metabolites were extracted using 400 μl ACN:H_2_O:HAc = 79:20:1 per 100 mg of the sample and incubation at 20°C, 180 rpm for 1 hour. The samples were analyzed by LC-MS/MS ^83^.

### RNA-seq (during infection)

For the RNA-seq experiment a type I inoculation was performed. 10 μl of a spore suspension (4*10^5^ spores/ml) were pipetted into each floret of the wheat ear (cv. Apogee) which was subsequently wrapped with a plastic bag containing 2 ml sterile water to maintain humidity. All plants were incubated at 20°C with day/night cycles 16h/8h. After 24 hours the plastic bags were removed. The ears were harvested after 2 and 4 days and 3 ears, respectively, were pooled for one sample. The ears were pulverized using Retsch mill (f = 30 * s^-1^, 30 sec.). RNA was extracted using Monarch® Total RNA Miniprep Kit according to the manufacturer’s instructions and sent to the sequencing facility.

RNA-seq data were aligned to the reference genome using STAR version 2.7.10a. FeatureCounts version 2.0.3 was used to get the counts of aligned reads in a genome annotation. For differential expression analysis DESeq2 version 1.44.0 was used

## Author contributions

TS: draft manuscript preparation, experimental design, phenotyping, Bioinformatic analysis, data evaluation; CZ: experimental design, strain preparation, phenotyping; FK: Extractions and analysis; MS: secondary metabolite analysis; JS: Experimental design, data evaluation, funding acquisition

## Contributions

TS: experimental design, phenotypic characterization, data evaluation, preparation of the manuscript; CZ: experimental design, phenotypic characterization; FK: RNA preparation, plant infection; MS: Toxin analysis, data evaluation; JS: experimental design, data evaluation, funding acquisition

**Supplementary Figure 1:**
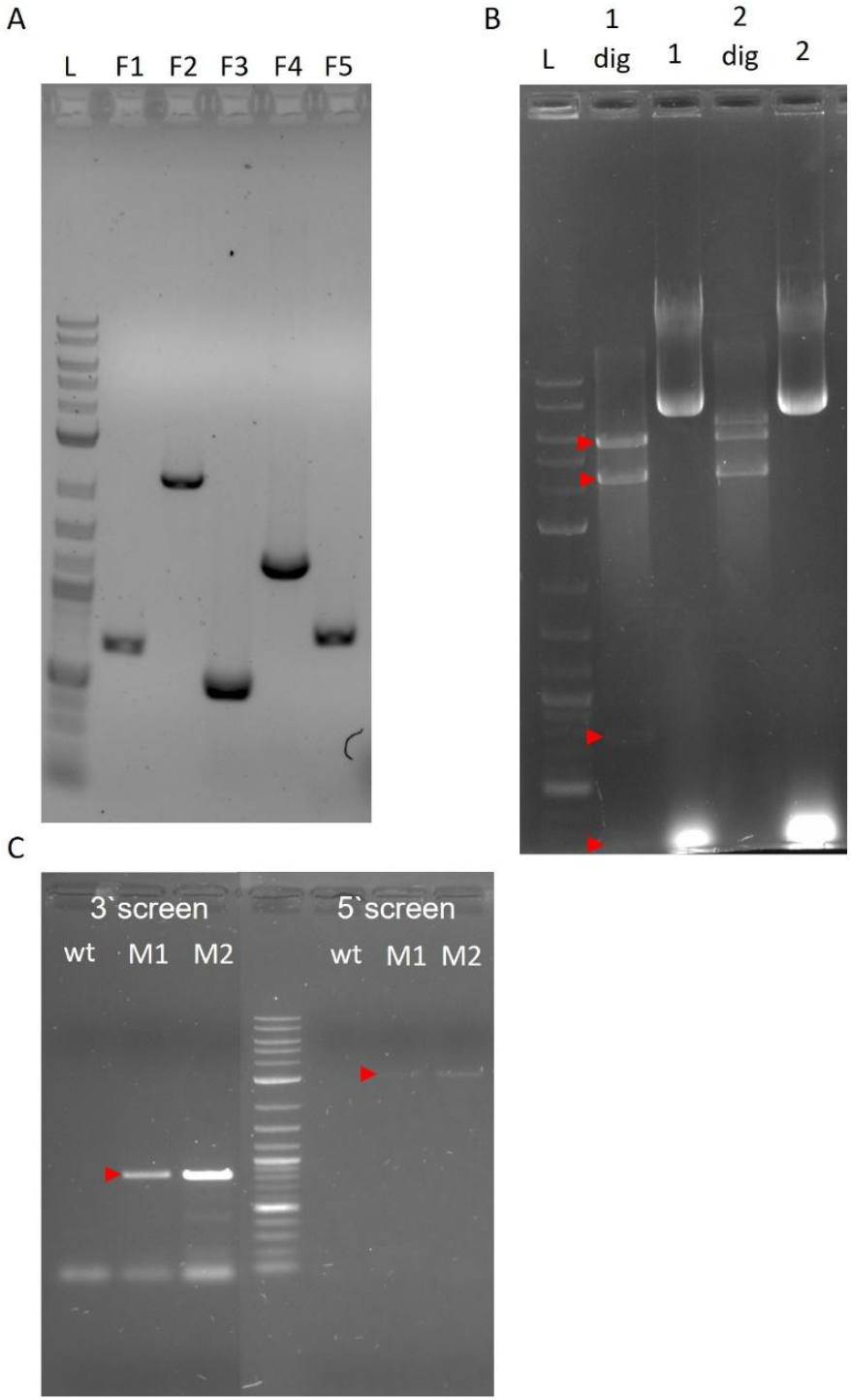
A: PCR fragments for plasmid assembly by yeast recombinational cloning; B: Test digest of two plasmid candidates used for introduction of the Ras-GAP mutation with EcoRI/BamHI. Expected size: 741 + 306 + 4286 + 5708 bp; C: diagnostic gel of PCR screening of transformed strains. Expected size: 3’mutant: 839 bp; 5’mutant: 3044 bp

**Supplementary Figure 2:**
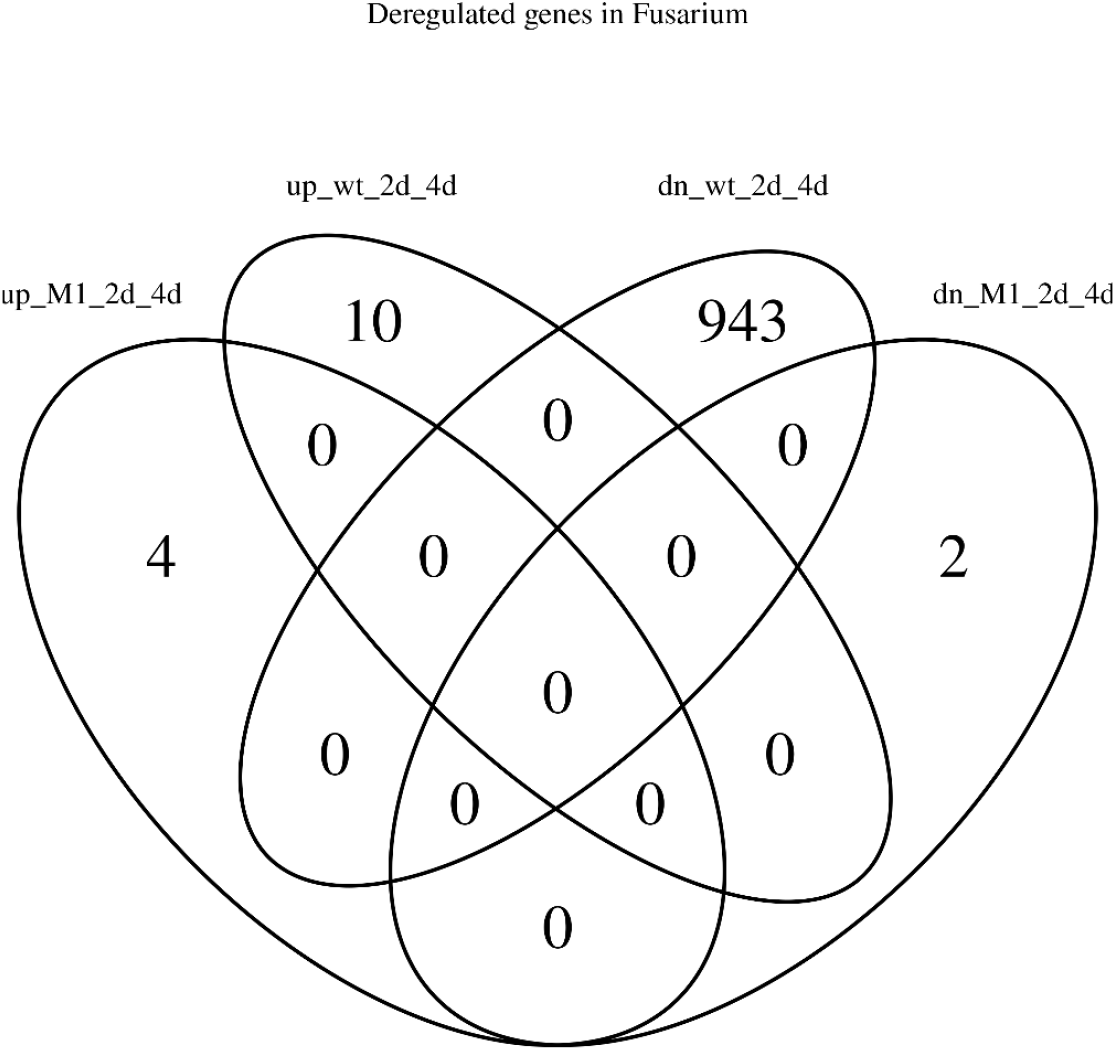
Venn diagram of genes significantly changing in the wild type and in the mutant between 2 dip and 4 dpi; up = upregulated, dn = down regulated, wt = wild type, M1 = Ras-GAP mutant strain

