## Supplementary File 1 for "Cell cycle controls pathogenic processes and mycotoxin production in *Fusarium graminearum*"

>FGSG_00355

MSSYDGSRSARQSKRYSMSALYMSISANEGDLQIEDELAKAQKSLRDLKSKISSQSKKNFVLEKDVRYLDSRIALLIQNRMALEEQNEVASHLEDALEMQQGGFPNDDKTQKYGNLLFLLQSEPRHIAHLCRLVSMSEIDSLLQTVMFTIYGNQYESREEHLLLTMFQSVLTYQFDNTPDYSSLLRANTPVSRMMTTYTRRGPGQSFLKSVLADRINGLIELKDLDLEINPLKVYERMIEQIEEDTGQLPPHLPKGITGEQAAENPQVQAIIEPRLTMLTEIANGFLTTIIEGLEEAPYGIRWICKQIRSLTKRKYPDANDQVICTLIGGFFFLRFINPAIVTPKSYMLIDGTPAERPRRTLTYIAKMLQNLANKPSYAKEPYMAKLQPFIHQNKDRINKFMIDLCEVQDFYESLEMDNYVALSKKDLELEITLNEVYAMHSLIDKHHSELWKDDNSHLAIIMSELGSSPPQLPRKENRVINLPLFSRWESAIGDLTAALDITQEEVYFMEAKSIFVQVMRSIPSTSGVARRPLRLERIADAAATNRSDAVMVRKGIRAMELLSQLQEMHVIDKTDQFSLLRDEVEQELQHLGSLKEGVIAETSKLQEVYKTIRDHNVYLNGQLETYKSYLHNVRSQSEGTKRKQQKQQVLGPYKFTHQQLEKEGVIQKSNVPDNRRANIYFNFTSPLPGTFVISLHYKGRNRGLLELDLKLDDLLEMQKDNQDDLDLEYVQFNVPKVLALLNKRFARKKGW

>MoSMO1

MSVMLQTPSRASSASSSSFQPISRQNTMSSYDGSRSTRQSKRYSMSALYMSMSANETDLEIEDDLAKAQKVLRDLKSKISSQSKKNFVLEKDVRYLDGRIALLIQNRMALEEQHETASYLEDAADLQEGAFPNDDRTQKYGNLMFLLQSEPRHIAHLCRLVSMQEIDSLLQTVMFTIYGNQYESREEHLLLTMFQSVLTYNFENTPEYSSLLRANTPVSRMMTTYTRRGPGQSFLKSVLADKINSLIELRDLDLEINPLKVYERMIEQIEEDTGSVPPSLPRGVTAEQAAANPQVQEIIEPRLAMLTDIANGFLSTIIDGLEEAPYGIRWICKQIRSLTKRKYPDANDQVICTLIGGFFFLRFINPAIVTPKSYMLIDGQPADKPRRTLTLIAKMLQNLANKPSYAKEPYMAGLQPFVQMNKSRVNKFMLDLCEVGDFYESLEMDNYVALSKKDLELSITLNEIYAMHGLLEKHAHELCRDENSHLALLMGELGQAPQQVPRKENRVMNLPLYSRWETAIDNLTAALDITQEEVFFMEAKSIFVQIMRSIPQNTAVARRPLRLERIADAAATSKNDAVMVRKGIRAMELLSQLQELKVTDKSDGFGLLRDEIEQELQHLGSLKEGVIAETQKLEEVYKTIRDHNAYLVGQLETYKSYLHNVRSQSEGTRRKAQKQQVLGPYKFTHQQLEKEGVIQKSNVPDNRRANIYFNFTSPLPGTFVISLHYKGRTRGLLELDLKLDDLLEMQKDGQDDLDLEYVQFNVTKVLALLNKRFARKKGW

>ScIra1

MNQSDPQDKKNFPMEYSLTKHLFFDRLLLVLPIESNLKTYADVEADSVFNSCRSIILNIAITKDLNPIIENTLGLIDLIVQDEEITSDNITDDIAHSILVLLRLLSDVFEYYWDQNNDFKKIRNDNYKPGFSSHRPNFHTSRPKHTRINPALATMLLCKISKLKFNTRTLKVLQNMSHHLSGSATISKSSILPDSQEFLQKRNYPAYTEKIDLTIDYIQRFISASNHVEFTKCVKTKVVAPLLISHTSTELGVVNHLDLFGCEYLTDKNLLAYLDILQHLSSYMKRTIFHSLLLYYASKAFLFWIMARPKEYVKIYNNLISSDYNSPSSSSDNGGSNNSDKTSISQLVSLLFDDVYSTFSVSSLLTNVNNDHHYHLHHSSSSSKTTNTNSPNSISKTSIKQSSVNASGNVSPSQFSTGNDASPTSPMASLSSPLNTNILGYPLSPITSTLGQANTSTSTTAATTKTDADTPSTMNTNNNNNNNNSANLNNIPQRIFSLDDISSFNSSRKSLNLDDSNSLFLWDTSQHSNASMTNTNMHAGVNNSQSQNDQSSLNYMENIMELYSNYTGSELSSHTAILRFLVVLTLLDSEVYDEMNSNSYRKISEPIMNINPKDSNTSSWGSASKNPSIRHLTHGLKKLTLQQGRKRNVKFLTYLIRNLNGGQFVSDVSLIDSIRSILFLMTMTSSISQIDSNIASVIFSKRFYNLLGQNLEVGTNWNSATANTFISHCVERNPLTHRRLQLEFFASGLQLDSDLFLRHLQLEKELNHIDLPKISLYTEGFRVFFHLVSTKKLHEDIAEKTSSVLKRLFCIIADILLKATPYFDDNVTKIIASILDGHILDQFDAARTLSNDDHVSFDAATSVYTEPTEIIHNSSDASLVSSLSQSPLSINSGSNITNTRTWDIQSILPTLSNRSSASDLSLSNILTNPLEAQQNNNANLLAHRLSGVPTTKRYASPNDSERSRQSPYSSPPQLQQSDLPSPLSVLSSSAGFSSNHSITATPTILKNIKSPKPNKTKKIADDKQLKQPSYSRVILSDNDEARKIMMNIFSIFKRMTNWFIRPDANTEFPKTFTDIIKPLFVSILDSNQRLQVTARAFIEIPLSYIATFEDIDNDLDPRVLNDHYLLCTYAVTLFASSLFDLKLENAKREMLLDIIVKFQRVRSYLSNLAEKHNLVQAIITTERLTLPLLVGAVGSGIFISLYCSRGNTPRLIKISCCEFLRSLRFYQKYVGALDQYSIYNIDFIDAMAQDNFTASGSVALQRRLRNNILTYIKGSDSILLDSMDVIYKKWFYFSCSKSVTQEELVDFRSLAGILASMSGILSDMQELEKSKSAPDNEGDSLSFESRNPAYEVHKSLKLELTKKMNFFISKQCQWLNNPNLLTRENSRDILSIELHPLSFNLLFNNLGLKIDELMSIDLSKSHEDSSFVLLEQIIIIIRTILKRDDDEKIMLLFSTDLLDAVDKLIEIVEKISIKSSKYYKGIIQMSKMFRAFEHSEKNLGISNHFHLKNKWLKLVIGWFKLSINKDYDFENLSRPLREMDLQKRDEDFLYIDTSIESAKALAYLTHNVPLEIPPSSSKEDWNRSSTVSFGNHFTILLKGLEKSADLNQFPVSLRHKISILNENVIIALTNLSNANVNVSLKFTLPMGYSPNKDIRIAFLRVFIDIVTNYPVNPEKHEMDKMLAIDDFLKYIIKNPILAFFGSLACSPADVDLYAGGFLNAFDTRNASHILVTELLKQEIKRAARSDDILRRNSCATRALSLYTRSRGNKYLIKTLRPVLQGIVDNKESFEIDKMKPGSENSEKMLDLFEKYMTRLIDAITSSIDDFPIELVDICKTIYNAASVNFPEYAYIAVGSFVFLRFIGPALVSPDSENIIIVTHAHDRKPFITLAKVIQSLANGRENIFKKDILVSKEEFLKTCSDKIFNFLSELCKIPTNNFTVNVREDPTPISFDYSFLHKFFYLNEFTIRKEIINESKLPGEFSFLKNTVMLNDKILGVLGQPSMEIKNEIPPFVVENREKYPSLYEFMSRYAFKKVDMKEEEEDNAPFVHEAMTLDGIQIIVVTFTNCEYNNFVMDSLVYKVLQIYARMWCSKHYVVIDCTTFYGGKANFQKLTTLFFSLIPEQASSNCMGCYYFNVNKSFMDQWASSYTVENPYLVTTIPRCFINSNTDQSLIKSLGLSGRSLEVLKDVRVTLHDITLYDKEKKKFCPVSLKIGNKYFQVLHEIPQLYKVTVSNRTFSIKFNNVYKISNLISVDVSNTTGVSSEFTLSLDNEEKLVFCSPKYLEIVKMFYYAQLKMEEDFGTDFSNDISFSTSSSAVNASYCNVKEVGEIISHLSLVILVGLFNEDDLVKNISYNLLVATQEAFNLDFGTRLHKSPETYVPDDTTTFLALIFKAFSESSTELTPYIWKYMLDGLENDVIPQEHIPTVVCSLSYWVPNLYEHVYLANDEEGPEAISRIIYSLIRLTVKEPNFTTAYLQQIWFLLALDGRLTNVIVEEIVSHALDRDSENRDWMKAVSILTSFPTTEIACQVIEKLINMIKSFLPSLAVEASAHSWSELTILSKISVSIFFESPLLSQMYLPEILFAVSLLIDVGPSEIRVSLYELLMNVCHSLTNNESLPERNRKNLDIVCATFARQKLNFISGFSQEKGRVLPNFAASSFSSKFGTLDLFTKNIMLLMEYGSISEGAQWEAKYKKYLMDAIFGHRSFFSARAMMILGIMSKSHTSLFLCKELLVETMKVFAEPVVDDEQMFIIIAHVFTYSKIVEGLDPSSELMKELFWLATICVESPHPLLFEGGLLFMVNCLKRLYTVHLQLGFDGKSLAKKLMESRNFAATLLAKLESYNGCIWNEDNFPHIILGFIANGLSIPVVKGAALDCLQALFKNTYYERKSNPKSSDYLCYLFLLHLVLSPEQLSTLLLEVGFEDELVPLNNTLKVPLTLINWLSSDSDKSNIVLYQGALLFSCVMSDEPCKFRFALLMRYLLKVNPICVFRFYTLTRKEFRRLSTLEQSSEAVAVSFELIGMLVTHSEFNYLEEFNDEMVELLKKRGLSVVKPLDIFDQEHIEKLKGEGEHQVAIYERKRLATMILARMSCS

>ScIra2

MSQPTKNKKKEHGTDSKSSRMTRTLVNHILFERILPILPVESNLSTYSEVEEYSSFISCRSVLINVTVSRDANAMVEGTLELIESLLQGHEIISDKGSSDVIESILIILRLLSDALEYNWQNQESLHYNDISTHVEHDQEQKYRPKLNSILPDYSSTHSNGNKHFFHQSKPQALIPELASKLLESCAKLKFNTRTLQILQNMISHVHGNILTTLSSSILPRHKSYLTRHNHPSHCKMIDSTLGHILRFVAASNPSEYFEFIRKSVQVPVTQTHTHSHSHSHSLPSSVYNSIVPHFDLFSFIYLSKHNFKKYLELIKNLSVTLRKTIYHCLLLHYSAKAIMFWIMARPAEYYELFNLLKDNNNEHSKSLNTLNHTLFEEIHSTFNVNSMITTNQNAHQGSSSPSSSSPSSPPSSSSSDNNNQNIIAKSLSRQLSHHQSYIQQQSERKLHSSWTTNSQSSTSLSSSTSNSTTTDFSTHTQPGEYDPSLPDTPTMSNITISASSLLSQTPTPTTQLQQRLNSAAAAAAAAASPSNSTPTGYTAEQQSRASYDAHKTGHTGKDYDEHFLSVTRLDNVLELYTHFDDTEVLPHTSVLKFLTTLTMFDIDLFNELNATSFKYIPDCTMHRPKERTSSFNNTAHETGSEKTSGIKHITQGLKKLTSLPSSTKKTVKFVKMLLRNLNGNQAVSDVALLDTMRALLSFFTMTSAVFLVDRNLPSVLFAKRLIPIMGTNLSVGQDWNSKINNSLMVCLKKNSTTFVQLQLIFFSSAIQFDHELLLARLSIDTMANNLNMQKLCLYTEGFRIFFDIPSKKELRKAIAVKISKFFKTLFSIIADILLQEFPYFDEQITDIVASILDGTIINEYGTKKHFKGSSPSLCSTTRSRSGSTSQSSMTPVSPLGLDTDICPMNTLSLVGSSTSRNSDNVNSLNSSPKNLSSDPYLSHLVAPRARHALGGPSSIIRNKIPTTLTSPPGTEKSSPVQRPQTESISATPMAITNSTPLSSAAFGIRSPLQKIRTRRYSDESLGKFMKSTNNYIQEHLIPKDLNEATLQDARRIMINIFSIFKRPNSYFIIPHNINSNLQWVSQDFRNIMKPIFVAIVSPDVDLQNTAQSFMDTLLSNVITYGESDENISIEGYHLLCSYTVTLFAMGLFDLKINNEKRQILLDITVKFMKVRSHLAGIAEASHHMEYISDSEKLTFPLIMGTVGRALFVSLYSSQQKIEKTLKIAYTEYLSAINFHERNIDDADKTWVHNIEFVEAMCHDNYTTSGSIAFQRRTRNNILRFATIPNAILLDSMRMIYKKWHTYTHSKSLEKQERNDFRNFAGILASLSGILFINKKILQEMYPYLLDTVSELKKNIDSFISKQCQWLNYPDLLTRENSRDILSVELHPLSFNLLFNNLRLKLKELACSDLSIPENESSYVLLEQIIKMLRTILGRDDDNYVMMLFSTEIVDLIDLLTDEIKKIPAYCPKYLKAIIQMTKMFSALQHSEVNLGVKNHFHVKNKWLRQITDWFQVSIAREYDFENLSKPLKEMDLVKRDMDILYIDTAIEASTAIAYLTRHTFLEIPPAASDPELSRSRSVIFGFYFNILMKGLEKSSDRDNYPVFLRHKMSVLNDNVILSLTNLSNTNVDASLQFTLPMGYSGNRNIRNAFLEVFINIVTNYRTYTAKTDLGKLEAADKFLRYTIEHPQLSSFGAAVCPASDIDAYAAGLINAFETRNATHIVVAQLIKNEIEKSSRPTDILRRNSCATRSLSMLARSKGNEYLIRTLQPLLKKIIQNRDFFEIEKLKPEDSDAERQIELFVKYMNELLESISNSVSYFPPPLFYICQNIYKVACEKFPDHAIIAAGSFVFLRFFCPALVSPDSENIIDISHLSEKRTFISLAKVIQNIANGSENFSRWPALCSQKDFLKECSDRIFRFLAELCRTDRTIDIQVRTDPTPIAFDYQFLHSFVYLYGLEVRRNVLNEAKHDDGDIDGDDFYKTTFLLIDDVLGQLGQPKMEFSNEIPIYIREHMDDYPELYEFMNRHAFRNIETSTAYSPSVHESTSSEGIPIITLTMSNFSDRHVDIDTVAYKFLQIYARIWTTKHCLIIDCTEFDEGGLDMRKFISLVMGLLPEVAPKNCIGCYYFNVNETFMDNYGKCLDKDNVYVSSKIPHYFINSNSDEGLMKSVGITGQGLKVLQDIRVSLHDITLYDEKRNRFTPVSLKIGDIYFQVLHETPRQYKIRDMGTLFDVKFNDVYEISRIFEVHVSSITGVAAEFTVTFQDERRLIFSSPKYLEIVKMFYYAQIRLESEYEMDNNSSTSSPNSNNKDKQQKERTKLLCHLLLVSLIGLFDESKKMKNSSYNLIAATEASFGLNFGSHFHRSPEVYVPEDTTTFLGVIGKSLAESNPELTAYMFIYVLEALKNNVIPHVYIPHTICGLSYWIPNLYQHVYLADDEEGPENISHIFRILIRLSVRETDFKAVYMQYVWLLLLDDGRLTDIIVDEVINHALERDSENRDWKKTISLLTVLPTTEVANNIIQKILAKIRSFLPSLKLEAMTQSWSELTILVKISIHVFFETSLLVQMYLPEILFIVSLLIDVGPRELRSSLHQLLMNVCHSLAINSALPQDHRNNLDEISDIFAHQKVKFMFGFSEDKGRILQIFSASSFASKFNILDFFINNILLLMEYSSTYEANVWKTRYKKYVLESVFTSNSFLSARSIMIVGIMGKSYITEGLCKAMLIETMKVIAEPKITDEHLFLAISHIFTYSKIVEGLDPNLDLMKHLFWFSTLFLESRHPIIFEGALLFVSNCIRRLYMAQFENESETSLISTLLKGRKFAHTFLSKIENLSGIVWNEDNFTHILIFIINKGLSNPFIKSTAFDFLKMMFRNSYFEHQINQKSDHYLCYMFLLYFVLNCNQFEELLGDVDFEGEMVNIENKNTIPKILLEWLSSDNENANITLYQGAILFKCSVTDEPSRFRFALIIRHLLTKKPICALRFYSVIRNEIRKISAFEQNSDCVPLAFDILNLLVTHSESNSLEKLHEESIERLTKRGLSIVTSSGIFAKNSDMMIPLDVKPEDIYERKRIMTMILSRMSCSA

>AN4998

MSVATMLQPASRASTSSSSSFQPIARQNTMSSHDTRSLRQSKRMSVTALYLSMSAKDRDLEISDDLARAQKFLRELKSKISSQSKKNFVLEKDVRYLDSRIALLIQNRMALEEQNEVANRLDDTVDPQEGFFPNDEKTQKYGNLLFLLQTEPRHIAHLCRLVSMSEIDSLLQTVMFTIYGNQYESREEHLLLTMFQSVLTYQFDNTPEYSSLLRQNTPVSRMMTTYTRRGPGQSYLKQVLAEQINALIELRDVDLEINPLKVYETMVRDIEEETGSLPDHLPRGVTGEVAAENPQVQAIIAPRLKKLTEIANSFLTTIINSVNQAPYGIRWICKQIRSLSRRKYPDAHDQTICTLIGGFFFLRFINPAIVTPRSYMLIDATPTDKPRRTLTLIAKMLQNLANKPSYAKEPYMAKLQPFIQQNKERVNKFMLDLCEVQDFYESLEMDNYVALSKRDLELQITLNEMYATHALLEKHNAALAQDQHSHLQEILQELGPAPPQLPRKENRTITVPLFSRWETALDDLTSALDITQEEVFFMEAKSTFVQILRSLPPHSSVARRPLRLDRIAEAAATLKNDAVMVRKGIRTMELLSQLQEMGVIDRSDEFSLLRDEVEQELVHLGSLKEKVMEETRQLESVYATIRDHNAYLVGQLETYKSYLQNVRSQSEGKSRKTQKQQELGPYKFTHQQLEKEGVIHKSNVPENRRANIYFMFKSPLPGTFVISLHYKGRARGLLELDLKLDDLLEMQKDNLEDLDLEYVQFNVSKVLTLLNKRFSRKKGW

Sc = Saccharomyces cerevisiae

Co = Colletotrichum obiculare

Fg = Fusarium graminearum

Mo = Magnaporthe oryzae

Phylogenetic tree:

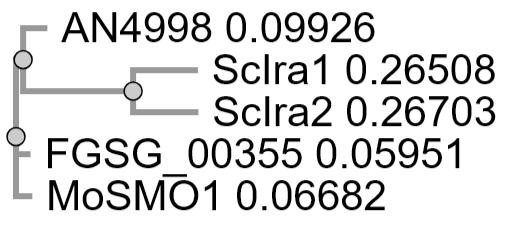

```
AN4998          ------------------------------------------------------------	0
FGSG_00355      ------------------------------------------------------------	0
MoSMO1          ------------------------------------------------------------	0
ScIra1          -------MNQSDPQDKKNFPMEYSLTKHLFFDRLLLVLPIESNLKTYADVEADSVFNSCR	53
ScIra2          MSQPTKNKKKEHGTDSKSSRMTRTLVNHILFERILPILPVESNLSTYSEVEEYSSFISCR	60


AN4998          ------------------------------------------------------------	0
FGSG_00355      ------------------------------------------------------------	0
MoSMO1          ------------------------------------------------------------	0
ScIra1          SIILNIAITKDLNPIIENTLGLIDLIVQDEEITSDNITDDIAHSILVLLRLLSDVFEYYW	113
ScIra2          SVLINVTVSRDANAMVEGTLELIESLLQGHEIISDKGSSDVIESILIILRLLSDALEYNW	120


AN4998          ------------------------------------------------------------	0
FGSG_00355      ------------------------------------------------------------	0
MoSMO1          ------------------------------------------------------------	0
ScIra1          DQNNDFK---------KIRNDNYKPGFSSHRPNFHTS----------RPKHTRINPALAT	154
ScIra2          QNQESLHYNDISTHVEHDQEQKYRPKLNSILPDYSSTHSNGNKHFFHQSKPQALIPELAS	180


AN4998          ------------------------------------------------------------	0
FGSG_00355      ------------------------------------------------------------	0
MoSMO1          ------------------------------------------------------------	0
ScIra1          MLLCKISKLKFNTRTLKVLQNMSHHLSGSAT-ISKSSILPDSQEFLQKRNYPAYTEKIDL	213
ScIra2          KLLESCAKLKFNTRTLQILQNMISHVHGNILTTLSSSILPRHKSYLTRHNHPSHCKMIDS	240


AN4998          ------------------------------------------------------------	0
FGSG_00355      ------------------------------------------------------------	0
MoSMO1          ------------------------------------------------------------	0
ScIra1          TIDYIQRFISASNHVEFTKCVKTKVVAPLLISHTS-----------TELGVVNHLDLFGC	262
ScIra2          TLGHILRFVAASNPSEYFEFIRKSVQVPVTQTHTHSHSHSHSLPSSVYNSIVPHFDLFSF	300


AN4998          ------------------------------------------------------------	0
FGSG_00355      ------------------------------------------------------------	0
MoSMO1          ------------------------------------------------------------	0
ScIra1          EYLTDKNLLAYLDILQHLSSYMKRTIFHSLLLYYASKAFLFWIMARPKEYVKIYNNLISS	322
ScIra2          IYLSKHNFKKYLELIKNLSVTLRKTIYHCLLLHYSAKAIMFWIMARPAEYYELFNLLKDN	360


AN4998          ------------------------------------------------------------	0
FGSG_00355      ------------------------------------------------------------	0
MoSMO1          ------------------------------------------------------------	0
ScIra1          DYNSPSSSSDNGGSNNSDKTSISQLVSLLFDDVYSTFSVSSLLTNVNNDHHYHLHHSSSS	382
ScIra2          N--------------NEHSKSLNTLNHTLFEEIHSTFNVNSMITTNQNAHQGSSSPSSSS	406


AN4998          ------------------------------------------------------------	0
FGSG_00355      ------------------------------------------------------------	0
MoSMO1          ------------------------------------------------------------	0
ScIra1          SKTT------NTNSPNSISKTSIKQSSVNA-----S-GNVSPSQFSTGNDASPTSPMASL	430
ScIra2          PSSPPSSSSSDNNNQNIIAKSLSRQLSHHQSYIQQQSERKLHSSWTTNSQS-STSLSSST	465


AN4998          ------------------------------------------------------------	0
FGSG_00355      ------------------------------------------------------------	0
MoSMO1          ------------------------------------------------------------	0
ScIra1          SSPLNTNILGYPLSPITSTLGQANTSTSTTAATTKTDADTPSTMNTNNNNNNNNSANLNN	490
ScIra2          ---SNSTTTDF----------------STHTQPGEYDPSLPDTPTMSN-ITISASSLLSQ	505


AN4998          ------------------------------------------------------------	0
FGSG_00355      ------------------------------------------------------------	0
MoSMO1          ------------------------------------------------------------	0
ScIra1          IPQRIFSLDDISSFNSSRKS--LNLDDSN-SLFLWDTSQHSNASMTNTNMHAGVNNSQSQ	547
ScIra2          TPTPTTQLQQ--RLNSAAAAAAAAASPSNSTPTGYTAEQQSRASYD--AHKTGHTGKDYD	561


AN4998          ------------------------------------------------------------	0
FGSG_00355      ------------------------------------------------------------	0
MoSMO1          ------------------------------------------------------------	0
ScIra1          NDQSSLNYMENIMELYSNYTGSELSSHTAILRFLVVLTLLDSEVYDEMNSNSYRKISEPI	607
ScIra2          EHFLSVTRLDNVLELYTHFDDTEVLPHTSVLKFLTTLTMFDIDLFNELNATSFKYIPDCT	621


AN4998          ------------------------------------------------------------	0
FGSG_00355      ------------------------------------------------------------	0
MoSMO1          ------------------------------------------------------------	0
ScIra1          MNINPKDS----NTSSWGSASKNPSIRHLTHGLKKLTL-QQGRKRNVKFLTYLIRNLNGG	662
ScIra2          MHRPKERTSSFNNTAHETGSEKTSGIKHITQGLKKLTSLPSSTKKTVKFVKMLLRNLNGN	681


AN4998          ------------------------------------------------------------	0
FGSG_00355      ------------------------------------------------------------	0
MoSMO1          ------------------------------------------------------------	0
ScIra1          QFVSDVSLIDSIRSILFLMTMTSSISQIDSNIASVIFSKRFYNLLGQNLEVGTNWNSATA	722
ScIra2          QAVSDVALLDTMRALLSFFTMTSAVFLVDRNLPSVLFAKRLIPIMGTNLSVGQDWNSKIN	741


AN4998          ------------------------------------------------------------	0
FGSG_00355      ------------------------------------------------------------	0
MoSMO1          ------------------------------------------------------------	0
ScIra1          NTFISHCVERNPLTHRRLQLEFFASGLQLDSDLFLRHLQLEKELNHIDLPKISLYTEGFR	782
ScIra2          NS-LMVCLKKNSTTFVQLQLIFFSSAIQFDHELLLARLSIDTMANNLNMQKLCLYTEGFR	800


AN4998          ------------------------------------------------------------	0
FGSG_00355      ------------------------------------------------------------	0
MoSMO1          ------------------------------------------------------------	0
ScIra1          VFFHLVSTKKLHEDIAEKTSSVLKRLFCIIADILLKATPYFDDNVTKIIASILDGHILDQ	842
ScIra2          IFFDIPSKKELRKAIAVKISKFFKTLFSIIADILLQEFPYFDEQITDIVASILDGTIINE	860


AN4998          ------------------------------------------------------------	0
FGSG_00355      ------------------------------------------------------------	0
MoSMO1          ------------------------------------------------------------	0
ScIra1          FDAARTLSNDDHVSFDAATSVYTEPTEIIHNSSDASLVSSLSQSPLSINSGSNITNTRTW	902
ScIra2          YGTKKHFKGSSPSLCSTTRSR--------SGS--TSQSSMTPVSPLGLDTDICPMN--TL	908


AN4998          ------------------------------------------------------------	0
FGSG_00355      ------------------------------------------------------------	0
MoSMO1          ------------------------------------------------------------	0
ScIra1          DIQSILPTLSNRSSAS----DLSLSNILTNPLEAQQNNNANLLAHRLSGVPTTKRY----	954
ScIra2          ---SLVGSSTSRNSDNVNSLNSSPKNLSSDPYLSHL--VAPRARHALGGPSSIIRNKIPT	963


AN4998          ------------------------------------------------------------	0
FGSG_00355      ------------------------------------------------------------	0
MoSMO1          ------------------------------------------------------------	0
ScIra1          --ASPNDSERSRQSPYSSPPQLQQSDLPSPLSVLSSSAGFSSNHSITATPTILKNIKSPK	1012
ScIra2          TLTSPPGTEK--SSPVQRPQTESISA--TPMAITNST-PLSS---------AAFGIRSPL	1009


AN4998          ------------------------------------------------------------	0
FGSG_00355      ------------------------------------------------------------	0
MoSMO1          ------------------------------------------------------------	0
ScIra1          PNK-TKKIADDK---QLK-QPSY-------SRVILSDNDEARKIMMNIFSIFKRMTNWFI	1060
ScIra2          QKIRTRRYSDESLGKFMKSTNNYIQEHLIPKDLNEATLQDARRIMINIFSIFKRPNSYFI	1069


AN4998          ------------------------------------------------------------	0
FGSG_00355      ------------------------------------------------------------	0
MoSMO1          ------------------------------------------------------------	0
ScIra1          RPDANT----EFPKTFTDIIKPLFVSILDSNQRLQVTARAFIEIPLSYIATFEDIDNDLD	1116
ScIra2          IPHNINSNLQWVSQDFRNIMKPIFVAIVSPDVDLQNTAQSFMDTLLSNVITYGESDENIS	1129


AN4998          ------------------------------------------------------------	0
FGSG_00355      ------------------------------------------------------------	0
MoSMO1          ------------------------------------------------------------	0
ScIra1          PRVLNDHYLLCTYAVTLFASSLFDLKLENAKREMLLDIIVKFQRVRSYLSNLAEKHNLVQ	1176
ScIra2          ---IEGYHLLCSYTVTLFAMGLFDLKINNEKRQILLDITVKFMKVRSHLAGIAEASHHME	1186


AN4998          ------------------------------------------------------------	0
FGSG_00355      ------------------------------------------------------------	0
MoSMO1          ------------------------------------------------------------	0
ScIra1          AIITTERLTLPLLVGAVGSGIFISLYCSRGNTPRLIKISCCEFLRSLRFYQKYVGALDQY	1236
ScIra2          YISDSEKLTFPLIMGTVGRALFVSLYSSQQKIEKTLKIAYTEYLSAINFHERNIDDADKT	1246


AN4998          ------------------------------------------------------------	0
FGSG_00355      ------------------------------------------------------------	0
MoSMO1          ------------------------------------------------------------	0
ScIra1          SIYNIDFIDAMAQDNFTASGSVALQRRLRNNILTYIKGSDSILLDSMDVIYKKWFYFSCS	1296
ScIra2          WVHNIEFVEAMCHDNYTTSGSIAFQRRTRNNILRFATIPNAILLDSMRMIYKKWHTYTHS	1306


AN4998          ------------------------------------------------------------	0
FGSG_00355      ------------------------------------------------------------	0
MoSMO1          ------------------------------------------------------------	0
ScIra1          KSVTQEELVDFRSLAGILASMSGILSDMQELEKSKSAPDNEGDSLSFESRNPAYEVHKSL	1356
ScIra2          KSLEKQERNDFRNFAGILASLSGILFINKKILQ------------------EMYPYLLDT	1348


AN4998          ------------------------------------------------------------	0
FGSG_00355      ------------------------------------------------------------	0
MoSMO1          ------------------------------------------------------------	0
ScIra1          KLELTKKMNFFISKQCQWLNNPNLLTRENSRDILSIELHPLSFNLLFNNLGLKIDELMSI	1416
ScIra2          VSELKKNIDSFISKQCQWLNYPDLLTRENSRDILSVELHPLSFNLLFNNLRLKLKELACS	1408


AN4998          ----------------------------------MSV--------ATMLQPASRASTSSS	18
FGSG_00355      ------------------------------------------------------------	0
MoSMO1          ------------------------------------M--------SVMLQTPSRASSASS	16
ScIra1          DLSKSHEDSSFVLLEQIIIIIRTILKRDDDEKIMLLFSTDLLDAVDKLIEIVEKISIKSS	1476
ScIra2          DLSIPENESSYVLLEQIIKMLRTILGRDDDNYVMMLFSTEIVDLIDLLTDEIKKIPAYCP	1468


AN4998          SSFQPIARQNTMSSHD-TRSLRQSKRMSVTALYLSMSAKDRDLEISDDLARAQKFLRELK	77
FGSG_00355      -----------MSSYDGSRSARQSKRYSMSALYMSISANEGDLQIEDELAKAQKSLRDLK	49
MoSMO1          SSFQPISRQNTMSSYDGSRSTRQSKRYSMSALYMSMSANETDLEIEDDLAKAQKVLRDLK	76
ScIra1          KYYKGI--------------------IQMSKMFRAFEHSEKNLGISNHFHLKNKWLKLVI	1516
ScIra2          KYLKAI--------------------IQMTKMFSALQHSEVNLGVKNHFHVKNKWLRQIT	1508
                                           .:: :: ::. .: :* :.:.:   :* *: :

AN4998          SKISSQSKKNFVL---------------EKDVRYLDSRI---------------------	101
FGSG_00355      SKISSQSKKNFVL---------------EKDVRYLDSRI---------------------	73
MoSMO1          SKISSQSKKNFVL---------------EKDVRYLDGRI---------------------	100
ScIra1          GWFKLSINKDYDFENLSRPLREMDLQKRDEDFLYIDTSIESAKALAYLTHNVPLEIPPSS	1576
ScIra2          DWFQVSIAREYDFENLSKPLKEMDLVKRDMDILYIDTAIEASTAIAYLTRHTFLEIPPAA	1568
                . :. .  ::: :               : *. *:*  *

AN4998          -----------------------------------------ALLIQNRM-ALEEQNEVAN	119
FGSG_00355      -----------------------------------------ALLIQNRM-ALEEQNEVAS	91
MoSMO1          -----------------------------------------ALLIQNRM-ALEEQHETAS	118
ScIra1          SKEDWNRSSTVSFGNHFTILLKGLEKSADLNQFPVSLRHKISILNENVIIALTNLSNANV	1636
ScIra2          SDPELSRSRSVIFGFYFNILMKGLEKSSDRDNYPVFLRHKMSVLNDNVILSLTNLSNTNV	1628
                                                         ::* :* : :* :  :.

AN4998          RLDDTVDPQEGFFPNDEKTQKYGNLLFLLQTEPRHIAHLCRLVSMSEIDSLLQTV----M	175
FGSG_00355      HLEDALEMQQGGFPNDDKTQKYGNLLFLLQSEPRHIAHLCRLVSMSEIDSLLQTV----M	147
MoSMO1          YLEDAADLQEGAFPNDDRTQKYGNLMFLLQSEPRHIAHLCRLVSMQEIDSLLQTV----M	174
ScIra1          NVSLKFTLPMGYSPNKDIRIAFLRVFIDIVTNYPVNPEKHEMDKMLAIDDFLKYIIKNPI	1696
ScIra2          DASLQFTLPMGYSGNRNIRNAFLEVFINIVTNYRTYTAKTDLGKLEAADKFLRYTIEHPQ	1688
                  .       *   * :    : .::: : ::         : .:   *.:*:

AN4998          FT-----------------IYGNQYESREE-HLLLTMFQSVLTYQFDNTPEYSSLLRQNT	217
FGSG_00355      FT-----------------IYGNQYESREE-HLLLTMFQSVLTYQFDNTPDYSSLLRANT	189
MoSMO1          FT-----------------IYGNQYESREE-HLLLTMFQSVLTYNFENTPEYSSLLRANT	216
ScIra1          LAFFGSLACSPADVDLYAGGFLNAFDTRNASHILV---TELLKQEIKRAARSDDILRRNS	1753
ScIra2          LSSFGAAVCPASDIDAYAAGLINAFETRNATHIVV---AQLIKNEIEKSSRPTDILRRNS	1745
                ::                    * :::*:  *:::    .::. ::..:    .:** *:

AN4998          PVSRMMTTYTRRGPGQSYLKQVLAEQINALIELRDVDLEINPLKVYETMVRDIEEETGSL	277
FGSG_00355      PVSRMMTTYTRRGPGQSFLKSVLADRINGLIELKDLDLEINPLKVYERMIEQIEEDTGQL	249
MoSMO1          PVSRMMTTYTRRGPGQSFLKSVLADKINSLIELRDLDLEINPLKVYERMIEQIEEDTGSV	276
ScIra1          CATRALSLYTRS-RGNKYLIKTLRPVLQGIVDNKES-FEIDKMKP---------------	1796
ScIra2          CATRSLSMLARS-KGNEYLIRTLQPLLKKIIQNRDF-FEIEKLKP---------------	1788
                 .:* ::  :*   *:.:*  .*   :: ::: ::  :**: :*

AN4998          PDHLPRGVTGEVAAENPQVQAIIAPRLKKLTEIANSFLTTIINSVNQAPYGIRWICKQIR	337
FGSG_00355      PPHLPKGITGEQAAENPQVQAIIEPRLTMLTEIANGFLTTIIEGLEEAPYGIRWICKQIR	309
MoSMO1          PPSLPRGVTAEQAAANPQVQEIIEPRLAMLTDIANGFLSTIIDGLEEAPYGIRWICKQIR	336
ScIra1          -----G-------------SENSEKMLDLFEKYMTRLIDAITSSIDDFPIELVDICKTIY	1838
ScIra2          -----E-------------DSDAERQIELFVKYMNELLESISNSVSYFPPPLFYICQNIY	1830
                                   .      :  : .  . :: :* ..:.  *  :  **: *

AN4998          SLSRRKYPDAHDQTICTLIGGFFFLRFINPAIVTPRSYMLIDATPTDKPRRTLTLIAKML	397
FGSG_00355      SLTKRKYPDANDQVICTLIGGFFFLRFINPAIVTPKSYMLIDGTPAERPRRTLTYIAKML	369
MoSMO1          SLTKRKYPDANDQVICTLIGGFFFLRFINPAIVTPKSYMLIDGQPADKPRRTLTLIAKML	396
ScIra1          NAASVNFPEYA----YIAVGSFVFLRFIGPALVSPDSENIIIVTHAHD-RKPFITLAKVI	1893
ScIra2          KVACEKFPDHA----IIAAGSFVFLRFFCPALVSPDSENIIDISHLSE-KRTFISLAKVI	1885
                . :  ::*:          *.*.****: **:*:* *  :*        :: :  :**::

AN4998          QNLANKP-SYAKEPYMAKLQPFIQQNKERVNKFMLDLCEVQDFYESLEMDNYV---ALSK	453
FGSG_00355      QNLANKP-SYAKEPYMAKLQPFIHQNKDRINKFMIDLCEVQDFYESLEMDNYV---ALSK	425
MoSMO1          QNLANKP-SYAKEPYMAGLQPFVQMNKSRVNKFMLDLCEVGDFYESLEMDNYV---ALSK	452
ScIra1          QSLANGRENIFKKDILVSKEEFLKTCSDKIFNFLSELCKIPTNNFTVNVREDPTPISFDY	1953
ScIra2          QNIANGSENFSRWPALCSQKDFLKECSDRIFRFLAELCRTD-RTIDIQVRTDPTPIAFDY	1944
                *.:**   .  :   :   : *::  ..:: .*: :**.       :::       ::.

AN4998          RDLELQITLNEMYATHALLEKH--------NAALAQDQHSHLQEILQELGPAPPQLPRKE	505
FGSG_00355      KDLELEITLNEVYAMHSLIDKH--------HSELWKDDNSHLAIIMSELGSSPPQLPRKE	477
MoSMO1          KDLELSITLNEIYAMHGLLEKH--------AHELCRDENSHLALLMGELGQAPQQVPRKE	504
ScIra1          SFLHKFFYLNEFTIRKEIINESKLPG----EFSFLKNTVMLNDKILGVLGQPSMEIKN--	2007
ScIra2          QFLHSFVYLYGLEVRRNVLNEAKHDDGDIDGDDFYKTTFLLIDDVLGQLGQPKMEFSN--	2002
                  *.  . *  .   : ::::            : :        ::  **    :. .

AN4998          NRTITVPLFSRWETALDDLTSALDITQEEVFFMEAKSTFVQILRSL--PPHSSVARRPLR	563
FGSG_00355      NRVINLPLFSRWESAIGDLTAALDITQEEVYFMEAKSIFVQVMRSI--PSTSGVARRPLR	535
MoSMO1          NRVMNLPLYSRWETAIDNLTAALDITQEEVFFMEAKSIFVQIMRSI--PQNTAVARRPLR	562
ScIra1          ----EIPPFVVEN--REKYPSL-------YEFMS-RYAFKKVDMKEEEEDNAPFVHEAMT	2053
ScIra2          ----EIPIYIREH--MDDYPEL-------YEFMN-RHAFRNIETS---TAYSPSVHESTS	2045
                     :* :   .    .             **. :  * ::  .      :  .:.

AN4998          LDRIAEAAATLK----NDAVMVRKGIRTMELLSQ---LQEMGVIDRSDEFSLLRDEVEQE	616
FGSG_00355      LERIADAAATNR----SDAVMVRKGIRAMELLSQ---LQEMHVIDKTDQFSLLRDEVEQE	588
MoSMO1          LERIADAAATSK----NDAVMVRKGIRAMELLSQ---LQELKVTDKSDGFGLLRDEIEQE	615
ScIra1          LDGIQIIVVTFTNCEYNNFVMDSLVYKVLQIYARMWCSKHYVVIDCTTF-----YGGKAN	2108
ScIra2          SEGIPIITLTMSNFSDRHVDIDTVAYKFLQIYARIWTTKHCLIIDCTEF-----DEGGLD	2100
                 : *   . *       .  :     : ::: ::    :.  : * :            :

AN4998          LVHLGSLKEKVMEETRQ---------------LESVYATIRDHNAYLVGQLETYKSYLQN	661
FGSG_00355      LQHLGSLKEGVIAETSK---------------LQEVYKTIRDHNVYLNGQLETYKSYLHN	633
MoSMO1          LQHLGSLKEGVIAETQK---------------LEEVYKTIRDHNAYLVGQLETYKSYLHN	660
ScIra1          FQKLTTLFFSLIPEQASSNCMGCYYFNVNKSFMDQWASSYTVENPYLVTTIPRCFINSNT	2168
ScIra2          MRKFISLVMGLLPEVAPKNCIGCYYFNVNETFMDNYGKCLDKDNVYVSSKIPHYFINSNS	2160
                : :: :*   :: *                  ::.       .* *:   :       :.

AN4998          VRS--QSEGKSRKTQKQQELGPYKFTHQQLEKEGVI-HKSNVPENRRANIYFMFKSPLPG	718
FGSG_00355      VRS--QSEGTKRKQQKQQVLGPYKFTHQQLEKEGVI-QKSNVPDNRRANIYFNFTSPLPG	690
MoSMO1          VRS--QSEGTRRKAQKQQVLGPYKFTHQQLEKEGVI-QKSNVPDNRRANIYFNFTSPLPG	717
ScIra1          DQSLIKSLGL--SGRSLEVLKDVRVTLHDITLYDKEKKKFCPVSLKIGNKYFQVLHEIPQ	2226
ScIra2          DEGLMKSVGI--TGQGLKVLQDIRVSLHDITLYDEKRNRFTPVSLKIGDIYFQVLHETPR	2218
                 ..  :* *   . :  : *   :.: :::   .   ::    . : .: ** .    *

AN4998          TFVISLHYKGRARGLLELDLKLDDLLEM------QKDNLEDLDL---------EYVQFNV	763
FGSG_00355      TFVISLHYKGRNRGLLELDLKLDDLLEM------QKDNQDDLDL---------EYVQFNV	735
MoSMO1          TFVISLHYKGRTRGLLELDLKLDDLLEM------QKDGQDDLDL---------EYVQFNV	762
ScIra1          LYKVTVS------NR-TFSIKFNNVYKISNLISVDVSNTTGVSSEFTLSLDNEEKLVFCS	2279
ScIra2          QYKIRDM------GT-LFDVKFNDVYEISRIFEVHVSSITGVAAEFTVTFQDERRLIFSS	2271
                 : :         .   :.:*:::: ::      . ..  .:           . : *

AN4998          SKVLTLLNKRFSRKK----GW---------------------------------------	780
FGSG_00355      PKVLALLNKRFARKK----GW---------------------------------------	752
MoSMO1          TKVLALLNKRFARKK----GW---------------------------------------	779
ScIra1          PKYLEIVKMFYYAQLKMEEDFGTDFSNDISFSTSSSAVNASYCNVKEVGEIISHLSLVIL	2339
ScIra2          PKYLEIVKMFYYAQIRLESEYEMDNNSST----SSPNSNNKDKQQKERTKLLCHLLLVSL	2327
                 * * :::  :  :      :

AN4998          ------------------------------------------------------------	780
FGSG_00355      ------------------------------------------------------------	752
MoSMO1          ------------------------------------------------------------	779
ScIra1          VGLFNEDDLVKNISYNLLVATQEAFNLDFGTRLHKSPETYVPDDTTTFLALIFKAFSESS	2399
ScIra2          IGLFDESKKMKNSSYNLIAATEASFGLNFGSHFHRSPEVYVPEDTTTFLGVIGKSLAESN	2387


AN4998          ------------------------------------------------------------	780
FGSG_00355      ------------------------------------------------------------	752
MoSMO1          ------------------------------------------------------------	779
ScIra1          TELTPYIWKYMLDGLENDVIPQEHIPTVVCSLSYWVPNLYEHVYLANDEEGPEAISRIIY	2459
ScIra2          PELTAYMFIYVLEALKNNVIPHVYIPHTICGLSYWIPNLYQHVYLADDEEGPENISHIFR	2447


AN4998          ------------------------------------------------------------	780
FGSG_00355      ------------------------------------------------------------	752
MoSMO1          ------------------------------------------------------------	779
ScIra1          SLIRLTVKEPNFTTAYLQQIWFLLALDGRLTNVIVEEIVSHALDRDSENRDWMKAVSILT	2519
ScIra2          ILIRLSVRETDFKAVYMQYVWLLLLDDGRLTDIIVDEVINHALERDSENRDWKKTISLLT	2507


AN4998          ------------------------------------------------------------	780
FGSG_00355      ------------------------------------------------------------	752
MoSMO1          ------------------------------------------------------------	779
ScIra1          SFPTTEIACQVIEKLINMIKSFLPSLAVEASAHSWSELTILSKISVSIFFESPLLSQMYL	2579
ScIra2          VLPTTEVANNIIQKILAKIRSFLPSLKLEAMTQSWSELTILVKISIHVFFETSLLVQMYL	2567


AN4998          ------------------------------------------------------------	780
FGSG_00355      ------------------------------------------------------------	752
MoSMO1          ------------------------------------------------------------	779
ScIra1          PEILFAVSLLIDVGPSEIRVSLYELLMNVCHSLTNNESLPERNRKNLDIVCATFARQKLN	2639
ScIra2          PEILFIVSLLIDVGPRELRSSLHQLLMNVCHSLAINSALPQDHRNNLDEISDIFAHQKVK	2627


AN4998          ------------------------------------------------------------	780
FGSG_00355      ------------------------------------------------------------	752
MoSMO1          ------------------------------------------------------------	779
ScIra1          FISGFSQEKGRVLPNFAASSFSSKFGTLDLFTKNIMLLMEYGSISEGAQWEAKYKKYLMD	2699
ScIra2          FMFGFSEDKGRILQIFSASSFASKFNILDFFINNILLLMEYSSTYEANVWKTRYKKYVLE	2687


AN4998          ------------------------------------------------------------	780
FGSG_00355      ------------------------------------------------------------	752
MoSMO1          ------------------------------------------------------------	779
ScIra1          AIFGHRSFFSARAMMILGIMSKSHTSLFLCKELLVETMKVFAEPVVDDEQMFIIIAHVFT	2759
ScIra2          SVFTSNSFLSARSIMIVGIMGKSYITEGLCKAMLIETMKVIAEPKITDEHLFLAISHIFT	2747


AN4998          ------------------------------------------------------------	780
FGSG_00355      ------------------------------------------------------------	752
MoSMO1          ------------------------------------------------------------	779
ScIra1          YSKIVEGLDPSSELMKELFWLATICVESPHPLLFEGGLLFMVNCLKRLYTVHLQLGFDGK	2819
ScIra2          YSKIVEGLDPNLDLMKHLFWFSTLFLESRHPIIFEGALLFVSNCIRRLYMAQFENES-ET	2806


AN4998          ------------------------------------------------------------	780
FGSG_00355      ------------------------------------------------------------	752
MoSMO1          ------------------------------------------------------------	779
ScIra1          SLAKKLMESRNFAATLLAKLESYNGCIWNEDNFPHIILGFIANGLSIPVVKGAALDCLQA	2879
ScIra2          SLISTLLKGRKFAHTFLSKIENLSGIVWNEDNFTHILIFIINKGLSNPFIKSTAFDFLKM	2866


AN4998          ------------------------------------------------------------	780
FGSG_00355      ------------------------------------------------------------	752
MoSMO1          ------------------------------------------------------------	779
ScIra1          LFKNTYYERKSNPKSSDYLCYLFLLHLVLSPEQLSTLLLEVGFEDELVPLNNTLKVPLTL	2939
ScIra2          MFRNSYFEHQINQKSDHYLCYMFLLYFVLNCNQFEELLGDVDFEGEMVNIENKNTIPKIL	2926


AN4998          ------------------------------------------------------------	780
FGSG_00355      ------------------------------------------------------------	752
MoSMO1          ------------------------------------------------------------	779
ScIra1          INWLSSDSDKSNIVLYQGALLFSCVMSDEPCKFRFALLMRYLLKVNPICVFRFYTLTRKE	2999
ScIra2          LEWLSSDNENANITLYQGAILFKCSVTDEPSRFRFALIIRHLLTKKPICALRFYSVIRNE	2986


AN4998          ------------------------------------------------------------	780
FGSG_00355      ------------------------------------------------------------	752
MoSMO1          ------------------------------------------------------------	779
ScIra1          FRRLSTLEQSSEAVAVSFELIGMLVTHSEFNYLEEFNDEMVELLKKRGLSVVKPLDIFDQ	3059
ScIra2          IRKISAFEQNSDCVPLAFDILNLLVTHSESNSLEKLHEESIERLTKRGLSIVTSSGIFAK	3046


AN4998          ----------------------------------	780
FGSG_00355      ----------------------------------	752
MoSMO1          ----------------------------------	779
ScIra1          EHIEKLKGEGEHQVAIYERKRLATMILARMSCS-	3092
ScIra2          NSDMM-IPLDVKPEDIYERKRIMTMILSRMSCSA	3079
```

Alignment 2:

```
AN4998          MSVATMLQPASRASTSSSSSFQPIARQNTMSSHD-TRSLRQSKRMSVTALYLSMSAKDRD	59
FGSG_00355      -----------------------------MSSYDGSRSARQSKRYSMSALYMSISANEGD	31
MoSMO1          --MSVMLQTPSRASSASSSSFQPISRQNTMSSYDGSRSTRQSKRYSMSALYMSMSANETD	58
                                             ***:* :** ***** *::***:*:**:: *

AN4998          LEISDDLARAQKFLRELKSKISSQSKKNFVLEKDVRYLDSRIALLIQNRMALEEQNEVAN	119
FGSG_00355      LQIEDELAKAQKSLRDLKSKISSQSKKNFVLEKDVRYLDSRIALLIQNRMALEEQNEVAS	91
MoSMO1          LEIEDDLAKAQKVLRDLKSKISSQSKKNFVLEKDVRYLDGRIALLIQNRMALEEQHETAS	118
                *:*.*:**:*** **:***********************.***************:*.*.

AN4998          RLDDTVDPQEGFFPNDEKTQKYGNLLFLLQTEPRHIAHLCRLVSMSEIDSLLQTVMFTIY	179
FGSG_00355      HLEDALEMQQGGFPNDDKTQKYGNLLFLLQSEPRHIAHLCRLVSMSEIDSLLQTVMFTIY	151
MoSMO1          YLEDAADLQEGAFPNDDRTQKYGNLMFLLQSEPRHIAHLCRLVSMQEIDSLLQTVMFTIY	178
                 *:*: : *:* ****::*******:****:**************.**************

AN4998          GNQYESREEHLLLTMFQSVLTYQFDNTPEYSSLLRQNTPVSRMMTTYTRRGPGQSYLKQV	239
FGSG_00355      GNQYESREEHLLLTMFQSVLTYQFDNTPDYSSLLRANTPVSRMMTTYTRRGPGQSFLKSV	211
MoSMO1          GNQYESREEHLLLTMFQSVLTYNFENTPEYSSLLRANTPVSRMMTTYTRRGPGQSFLKSV	238
                **********************:*:***:****** *******************:**.*

AN4998          LAEQINALIELRDVDLEINPLKVYETMVRDIEEETGSLPDHLPRGVTGEVAAENPQVQAI	299
FGSG_00355      LADRINGLIELKDLDLEINPLKVYERMIEQIEEDTGQLPPHLPKGITGEQAAENPQVQAI	271
MoSMO1          LADKINSLIELRDLDLEINPLKVYERMIEQIEEDTGSVPPSLPRGVTAEQAAANPQVQEI	298
                **::**.****:*:*********** *:.:***:**.:*  **:*:*.* ** ***** *

AN4998          IAPRLKKLTEIANSFLTTIINSVNQAPYGIRWICKQIRSLSRRKYPDAHDQTICTLIGGF	359
FGSG_00355      IEPRLTMLTEIANGFLTTIIEGLEEAPYGIRWICKQIRSLTKRKYPDANDQVICTLIGGF	331
MoSMO1          IEPRLAMLTDIANGFLSTIIDGLEEAPYGIRWICKQIRSLTKRKYPDANDQVICTLIGGF	358
                * ***  **:***.**:***:.:::***************::******:**.********

AN4998          FFLRFINPAIVTPRSYMLIDATPTDKPRRTLTLIAKMLQNLANKPSYAKEPYMAKLQPFI	419
FGSG_00355      FFLRFINPAIVTPKSYMLIDGTPAERPRRTLTYIAKMLQNLANKPSYAKEPYMAKLQPFI	391
MoSMO1          FFLRFINPAIVTPKSYMLIDGQPADKPRRTLTLIAKMLQNLANKPSYAKEPYMAGLQPFV	418
                *************:******. *:::****** ********************* ****:

AN4998          QQNKERVNKFMLDLCEVQDFYESLEMDNYVALSKRDLELQITLNEMYATHALLEKHNAAL	479
FGSG_00355      HQNKDRINKFMIDLCEVQDFYESLEMDNYVALSKKDLELEITLNEVYAMHSLIDKHHSEL	451
MoSMO1          QMNKSRVNKFMLDLCEVGDFYESLEMDNYVALSKKDLELSITLNEIYAMHGLLEKHAHEL	478
                : **.*:****:***** ****************:****.*****:** *.*::**   *

AN4998          AQDQHSHLQEILQELGPAPPQLPRKENRTITVPLFSRWETALDDLTSALDITQEEVFFME	539
FGSG_00355      WKDDNSHLAIIMSELGSSPPQLPRKENRVINLPLFSRWESAIGDLTAALDITQEEVYFME	511
MoSMO1          CRDENSHLALLMGELGQAPQQVPRKENRVMNLPLYSRWETAIDNLTAALDITQEEVFFME	538
                 :*::***  :: *** :* *:******.:.:**:****:*:.:**:*********:***

AN4998          AKSTFVQILRSLPPHSSVARRPLRLDRIAEAAATLKNDAVMVRKGIRTMELLSQLQEMGV	599
FGSG_00355      AKSIFVQVMRSIPSTSGVARRPLRLERIADAAATNRSDAVMVRKGIRAMELLSQLQEMHV	571
MoSMO1          AKSIFVQIMRSIPQNTAVARRPLRLERIADAAATSKNDAVMVRKGIRAMELLSQLQELKV	598
                *** ***::**:*  :.********:***:**** :.**********:*********: *

AN4998          IDRSDEFSLLRDEVEQELVHLGSLKEKVMEETRQLESVYATIRDHNAYLVGQLETYKSYL	659
FGSG_00355      IDKTDQFSLLRDEVEQELQHLGSLKEGVIAETSKLQEVYKTIRDHNVYLNGQLETYKSYL	631
MoSMO1          TDKSDGFGLLRDEIEQELQHLGSLKEGVIAETQKLEEVYKTIRDHNAYLVGQLETYKSYL	658
                 *::* *.*****:**** ******* *: ** :*:.** ******.** **********

AN4998          QNVRSQSEGKSRKTQKQQELGPYKFTHQQLEKEGVIHKSNVPENRRANIYFMFKSPLPGT	719
FGSG_00355      HNVRSQSEGTKRKQQKQQVLGPYKFTHQQLEKEGVIQKSNVPDNRRANIYFNFTSPLPGT	691
MoSMO1          HNVRSQSEGTRRKAQKQQVLGPYKFTHQQLEKEGVIQKSNVPDNRRANIYFNFTSPLPGT	718
                :********. ** **** *****************:*****:******** *.******


AN4998          FVISLHYKGRARGLLELDLKLDDLLEMQKDNLEDLDLEYVQFNVSKVLTLLNKRFSRKKG	779
FGSG_00355      FVISLHYKGRNRGLLELDLKLDDLLEMQKDNQDDLDLEYVQFNVPKVLALLNKRFARKKG	751
MoSMO1          FVISLHYKGRTRGLLELDLKLDDLLEMQKDGQDDLDLEYVQFNVTKVLALLNKRFARKKG	778
                ********** *******************. :*********** ***:******:****

AN4998          W	780
FGSG_00355      W	752
MoSMO1          W	779
                *
```
